# Chemotherapy reshapes cellular clonal landscape via persister programs in breast cancer xenografts

**DOI:** 10.64898/2026.04.02.716097

**Authors:** Sarah Kronheim, Jiahui Wang, Jiao Yi Chen, Daniel Guerrero-Romero, Suet-Feung Chin, Giulia Lerda, Allan J.W. Lui, Oscar M. Rueda, Federico Gaiti, Carlos Caldas, Long V. Nguyen

## Abstract

Tumor ecosystems evolve in response to chemotherapy, but how treatment reshapes clonal architecture and the transcriptional programs of distinct cell states remains unclear. To investigate the response to chemotherapy by individual single cell-derived clones, we used a high-complexity lentiviral barcoding strategy by genetically labelling with unique, heritable and expressible barcode sequences individual cells from four human breast cancer patient-derived tumor xenograft models. We tracked 3,248 single cell-derived clones across 39 xenografts and profiled 676,292 cells with single cell RNA sequencing. The most striking changes following chemotherapy occurred in non-responsive xenografts, attributed to the emergence of previously minor cell clones that are chemotherapy resistant. In a triple negative model, epithelial and mesenchymal cell states showed distinct sensitivities to chemotherapy. ER+/HER2-models activated stress adaptive programs in response to carboplatin, highlighting DUSP1 and KLF4 as markers of platinum tolerance and a slow cycling, persister-like state. Together, these findings reveal that chemotherapy triggers an immediate and substantial reorganization of the cellular clonal landscape, driven by the selective engagement of stress response programs in cell clones that survive treatment.

## INTRODUCTION

Tumors are dynamic populations of cells capable of adaptation and selection under cytotoxic treatment pressure. These heterogeneous tumor cell populations differ in their genetic, transcriptional and phenotypic states, providing a substrate for differential responses to therapy. Pre-existing and adaptive tumor cell states can be differentially selected during therapy, reshaping the cellular clonal landscape of the tumor. Despite advances in systemic treatment, in particular targeted therapies and immune checkpoint inhibitors, chemotherapy remains a standard treatment for most cancers. Chemotherapy resistance therefore remains a major cause of treatment failure and disease progression.

Tracking the progeny of a single-cell using lentiviral-based genetic barcoding is a powerful approach for studying clonal cell lineage and its fitness and cell state heterogeneity. The progeny produced from a single uniquely genetically barcoded initiating cell^1–5^, is hereon referred to as a ‘cell clone’. The approach enables characterizing how distinct cell clones emerge, expand or decline within a complex heterogeneous tumor^6–9^. When combined with single-cell gene expression profiling this allows cell clone response to treatment to be directly associated with its transcriptional programs^10–13^. This combined approach enables identifying cell states that are primed to resist treatment and cell state transitions in response to treatment. Cell clones are distinct from genomic clones, and while there can be some overlap, the molecular and functional heterogeneity observed in the former is largely attributed to differences in transcriptional cell states that can be influenced by multiple factors, including the tumor microenvironment and epigenetic programming.

The concept of cancer persister cells are cells that survive therapeutic stress by entering reversible drug-tolerant states, rather than through permanent genomic alterations that confer treatment resistance^14,15^. It is not clear how chemotherapy resistance emerges in human breast cancer – if it is through persistence of pre-existing clones, adaptive cell state transitions induced by treatment-related stress, or a combination of both.

To assess this, we applied high-resolution lineage tracing in human breast cancer patient-derived tumor xenograft (PDTX) models treated with chemotherapy to investigate how the intratumoral cell clone composition shifts under therapeutic pressure. Coupling this approach with single cell RNA sequencing (scRNAseq) enabled us to directly link individual cell clones with their gene expression profiles, revealing transcriptomic features of clonal response that would not be feasible with bulk analysis. Chemotherapy induced marked and rapid shifts in the cellular clonal landscape, including the emergence of cell clones enriched for stress-adaptive gene programs. In a triple-negative model, epithelial and mesenchymal cell states showed distinct sensitivities to paclitaxel and carboplatin. In luminal B/ER+ tumors, we identify *DUSP1* and *KLF4* as candidate markers of platinum tolerance and a slow-cycling, persister-like state primed to withstand carboplatin exposure. Altogether, we find that in response to chemotherapy treatment there is an immediate and dramatic reconfiguration of the cellular clonal landscape with specific cell clones that modulate cellular stress responses to persist and survive despite treatment.

## RESULTS

### Xenografts established from single cell-derived clones reveal bulk and clonal response to treatment

To investigate response to chemotherapy on a cellular clonal level, we established genetically barcoded xenografts from four patient-derived tumor xenograft (PDTX) models (**Table S1 and Methods**). Barcoded xenografts were passaged into multiple secondary mice to establish cohorts in triplicate: untreated, paclitaxel-treated, and carboplatin-treated (**Fig. 1A**). This experimental design allowed for cell clones to propagate across multiple biological replicates and treatment cohorts, so the same cell clones can be directly compared with and without treatment.

**Figure 1.**
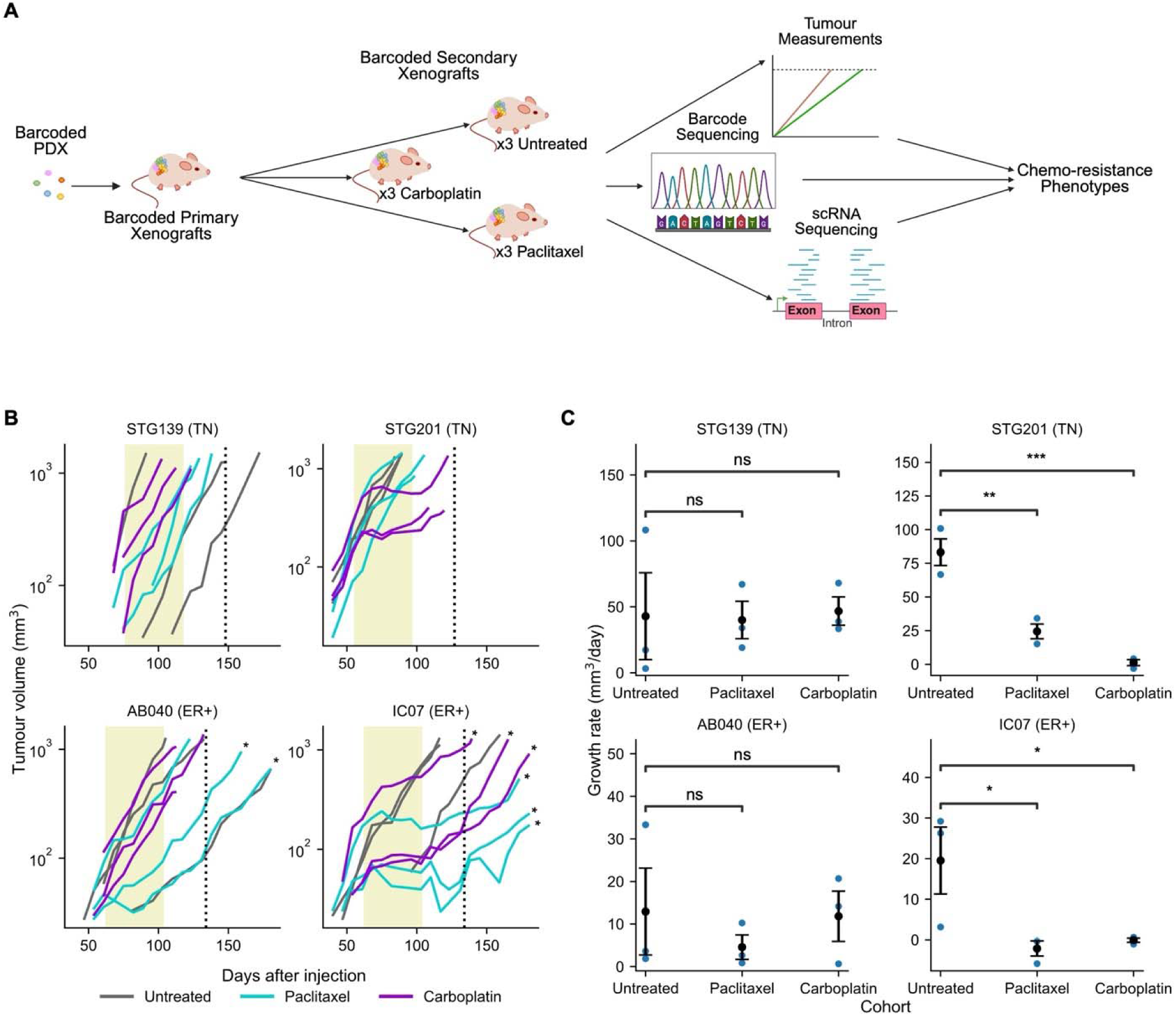
Overall experimental design and treatment response. (**A**) Experimental design. Three cohorts of secondary barcoded xenografts (untreated, carboplatin-treated, and paclitaxel-treated) were analyzed by tumour growth measurements, DNA amplicon barcode sequencing, and scRNAseq. (**B**) Tumour volume over time colored by cohort, * – >30 days of regrowth post-treatment before collection. (**C**) Bulk tumour growth rate during final week of treatment. Blue dots represent individual tumour measurements. Mean and standard error are represented with a black dot and error bars, respectively. One-tailed studenfs t-test shown as ns = not significant, * = p v 0.05, ** = p < 0.01, *** = _p_ < 0,001.

We then measured the change in bulk tumor growth rate with treatment to determine whether the xenograft model was responsive or non-responsive to carboplatin or paclitaxel chemotherapy. Mice were treated with chemotherapy for 6 weeks (corresponding to 2 cycles of treatment, **Fig. 1B**), and the tumor growth rate during the final week of treatment was assessed (**Table S2**). Overall, 1 triple-negative (TN) model STG139 and 1 ER+/HER2-(ER+) model AB040 were non-responsive to chemotherapy (absence of a statistically significant decrease in tumor growth rate compared to untreated control), and 1 TN model STG201 and 1 ER+ model IC07 were responsive to chemotherapy (presence of a statistically significant decrease in tumor growth rate compared to untreated control). STG201 demonstrated a 70% decrease in growth rate with paclitaxel and a 98% decrease in growth rate with carboplatin; IC07 demonstrated decreases in growth rate of 111% and 100%, respectively (**Fig. 1C**).

Using DNA amplicon sequencing we then quantified the number, frequency and type of tumor cell clones in each xenograft to map the changes in landscape with treatment. From analysis of 4 primary xenografts (1 per model), and 35 secondary xenografts from all 3 treatment cohorts, we detected a total of 3,248 single cell-derived clones (579 in primary and 2,722 in secondary xenografts) (**Fig. S1**). We did not observe consistent significant differences in the number of unique cell clones in secondary xenografts between treatment cohorts (**Fig. S2**).

Each single cell-derived clone was classified as either propagating (detected in both the primary and at least one secondary xenograft), transient (detected only in the primary but not secondary xenografts), or emerging (detected in at least one secondary but not the primary xenograft, **Table S3 and Fig. S3A**). Across all PDTX models, we detected 53/3,248 (1.6%) propagating, 526/3,248 (16%) transient, and 2,669/3,248 (82%) emerging single cell-derived clones. The frequency of propagating clones out of total barcoded cells used to establish the primary xenografts ranged from 1 in 120,000 to 1 in 22,000 cells, indicating they were rare, but their number and frequency did not consistently change with either paclitaxel or carboplatin treatment (**Fig. S3B**).

These data indicate that chemotherapy drives clonal remodeling primarily by enabling the emergence of previously silent cell populations, not by altering the pool of rare propagating clones. This pattern is consistent with the presence of dormant tumor cells that remain undetected at baseline but expand after treatment, even in xenografts that show no measurable response by tumor-volume criteria.

### Significant cellular clonal landscape changes identified in chemotherapy non-responsive xenografts

To further compare the changes in the landscape of cellular clonal composition in xenografts following chemotherapy, we classified cellular clonal response based on clone size determined by DNA amplicon sequencing. This classification was used to directly compare the size of clones irrespective of whether the clone was propagating or emerging. A cell clone was classified as “diminished” if its size in the treated xenograft was at least 30% smaller than its mean size in the untreated cohort, “expanded” if its size in the treated xenograft was at least 20% larger than its mean size in the untreated cohort, and “minimal change” in the remainder (**Methods**).

In models that were non-responsive to chemotherapy, we observed a rapid shift in cellular clonal composition in response to treatment. In TN model STG139, a single dominant (and propagating) cell clone in the untreated xenografts (where it represented 90-96% of barcoded cells) was significantly diminished following both paclitaxel treatment (representing 1-18% of barcoded cells) and carboplatin treatment (representing 0-2% of barcoded cells). It was replaced by 44-190 non-dominant emerging cell clones in paclitaxel-treated xenografts (representing 82-99% of barcoded cells) and 39-84 non-dominant emerging clones in carboplatin-treated xenografts (representing 98-100% of barcoded cells) (**Fig. 2A and Fig. S2B**). This shows a dramatic alteration of the cellular clonal landscape with chemotherapy in a model that based on bulk tumor measurements is non-responsive to treatment (**Fig. 2A**).

**Figure 2.**
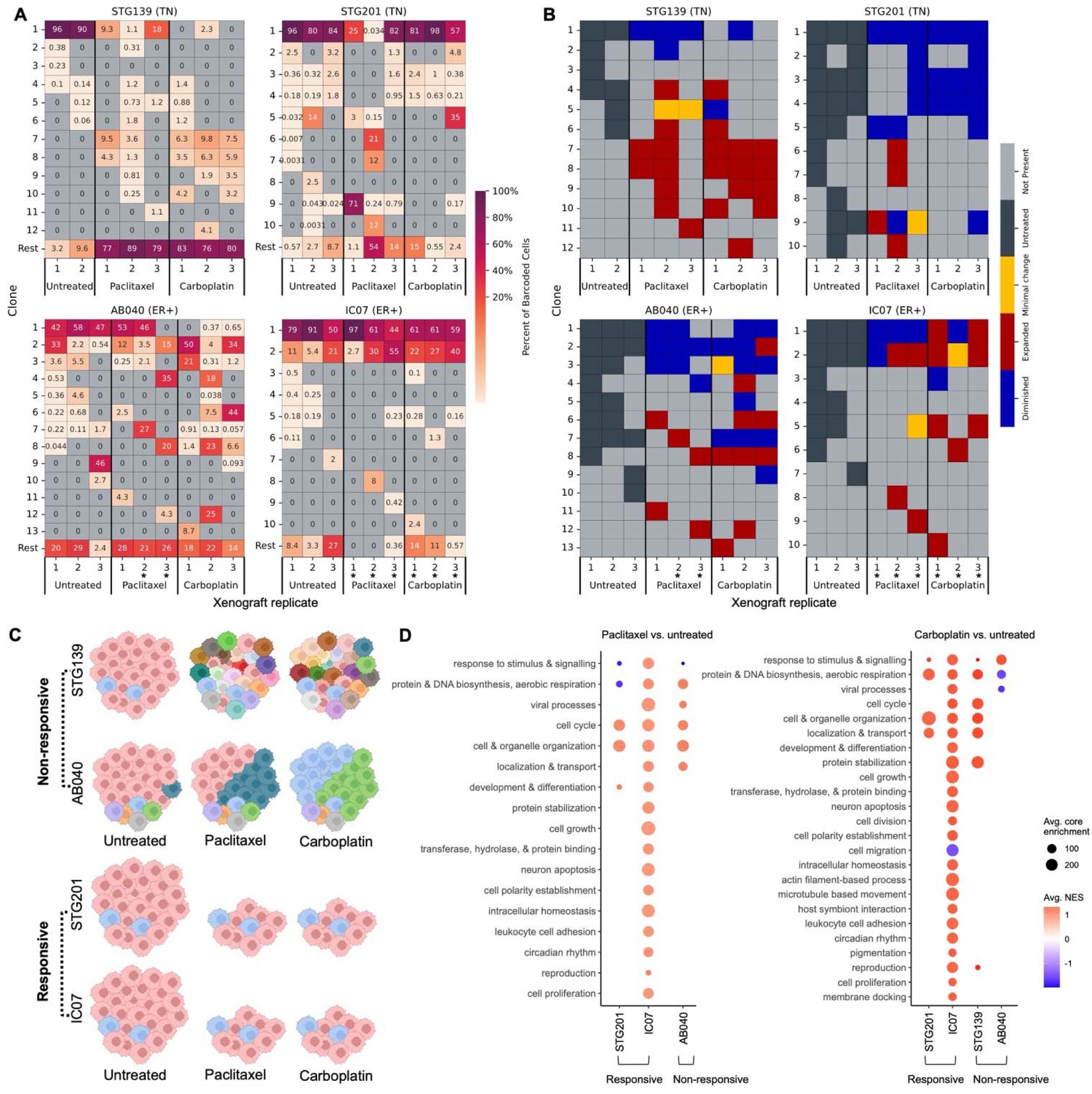
Rapid alteration of cellular clonal landscape and gene expression with chemotherapy. (**A**) and **(B)** Heatmap representing clone prevalence and response to treatment respectively per xenograft replicate. In (A), the bottom row groups the remaining clones together. Gray squares indicate absence of a clone, * = >30 days of regrowth post-treatment before tumor harvest. In (A) colours and numbers indicate the percent each clone contributes to the entire xenograft; in (B) colours indicate clonal response to treatment* (C) Model for clonal composition of overall sensitive and resistant PDTX with and without treatment. (D) Pathway expression differences on the cohort level comparing paclitaxel (left) and carboplatin (right) to untreated xenografts. TN model STG139 did not show significant pathway differences with paclitaxel treatment.

In ER+ model AB040, the two dominant (and propagating) cell clones in the untreated xenografts (defined as cell clones representing ≥20% of barcoded cells in at least one of the untreated xenografts) were variably responsive to chemotherapy. Interestingly, clones 4, 6, 8 and 12 were all emerging clones that were below the limit of detection in primary xenografts, were below the limit of detection or represented <1% of barcoded cells in all untreated xenografts and then emerged to become dominant cell clones representing up to 35%, 44%, 23% and 25% in individual chemotherapy-treated xenografts. In AB040, contrasting with STG139, there is a variable response to chemotherapy in the dominant cell clones, with examples of emerging clones that become dominant following chemotherapy treatment (**Fig. 2A-C**). These findings suggest an inherent biological difference in their response to chemotherapy. This is particularly striking because the tumor growth rates are unaffected by chemotherapy, and the observed significant alteration in cellular clonal representation would not be known or suspected without single-cell progeny clonal tracking analysis.

In sharp contrast, ER+ model IC07 with bulk tumor growth measurements that showed it was responsive to chemotherapy (**Fig. 1C**), had its cellular clonal representation largely unchanged. The two dominant (and propagating) cell clones in the untreated xenografts remained dominant following both paclitaxel treatment and carboplatin treatment (**Fig. 2A**). The most dominant cell clone was diminished with paclitaxel treatment but mostly expanded with carboplatin treatment, whereas the other dominant cell clone expanded in most xenografts in response to both chemotherapy agents **(Fig. 2B-C).**

The TN model STG201 was also responsive to chemotherapy. Here, the one dominant cell clone in the untreated xenografts diminished in size following both paclitaxel and carboplatin (**Fig. 2A**). But whereas this cell clone remained dominant following carboplatin treatment it variably diminished substantially following paclitaxel treatment and a new dominant cell clone appeared. For example, in one paclitaxel treated xenograft, a propagating cell clone represented 71% of barcoded cells and it represented only 0-0.03% of barcoded cells in the untreated cohort (and 10% of barcoded cells in the primary xenograft). In another paclitaxel treated xenograft, an emerging cell clone appears representing 21% of barcoded cells, when it represented <0.01% of cells in untreated xenografts and was not detected in the primary xenograft.

These data demonstrate that chemotherapy exerts substantial selective pressure on the cellular clonal architecture of these tumors, and that these changes are not reliably captured by bulk measurements of tumor response. Non-responsive xenografts show the greatest cellular clonal turnover, whereas responsive tumors remain dominated by cell clones that withstand treatment. These observations underscore the importance of cellular clonal-resolution analyses for interpreting therapeutic response.

### Single cell RNA sequencing identifies cell clusters associated with treatment response

Across four primary and 35 secondary barcoded xenografts from the 3 treatment cohorts, scRNAseq analysis yielded 676,292 high-quality single-cell transcriptomes (**Fig. S4A** and **Methods**). Differential gene expression comparing treated and untreated xenografts (**Fig. S5 and Table S4**) revealed broad transcriptional shifts following chemotherapy. Pathway enrichment analysis showed that paclitaxel and carboplatin activated programs associated with cell cycle progression, cytoskeletal and organelle organization, and intracellular transport – consistent with a proliferative rebound and clonal expansion after treatment (**Fig. 2D and S6**). Mitochondrial oxidative phosphorylation and aerobic respiration were also frequently upregulated; for example, the TN model STG139 exhibited marked induction of these pathways in response to carboplatin (**Fig. S6**). Notably, the ER+ model IC07 displayed the most significantly upregulated pathways, a finding that aligns with the extended interval between treatment and xenograft tumor tissue harvest in this model (**Fig. 1B**).

To define how chemotherapy reshapes transcriptional cell states, we examined the single-cell transcriptional profiles in greater detail. Each model resolved into 9-12 transcriptionally distinct cell clusters (**Fig. S4B**). Assessing transcriptional cell cluster composition based on contributions from treated versus untreated xenografts (**Fig. S7A**), we identified 1-3 transcriptional cell clusters per model that exhibited clear shifts in representation following treatment (**Fig. 3A**). These changes indicate a substantive reconfiguration of the cellular transcriptional landscape after chemotherapy, with specific cell states becoming either depleted or enriched under therapeutic pressure.

**Figure 3.**
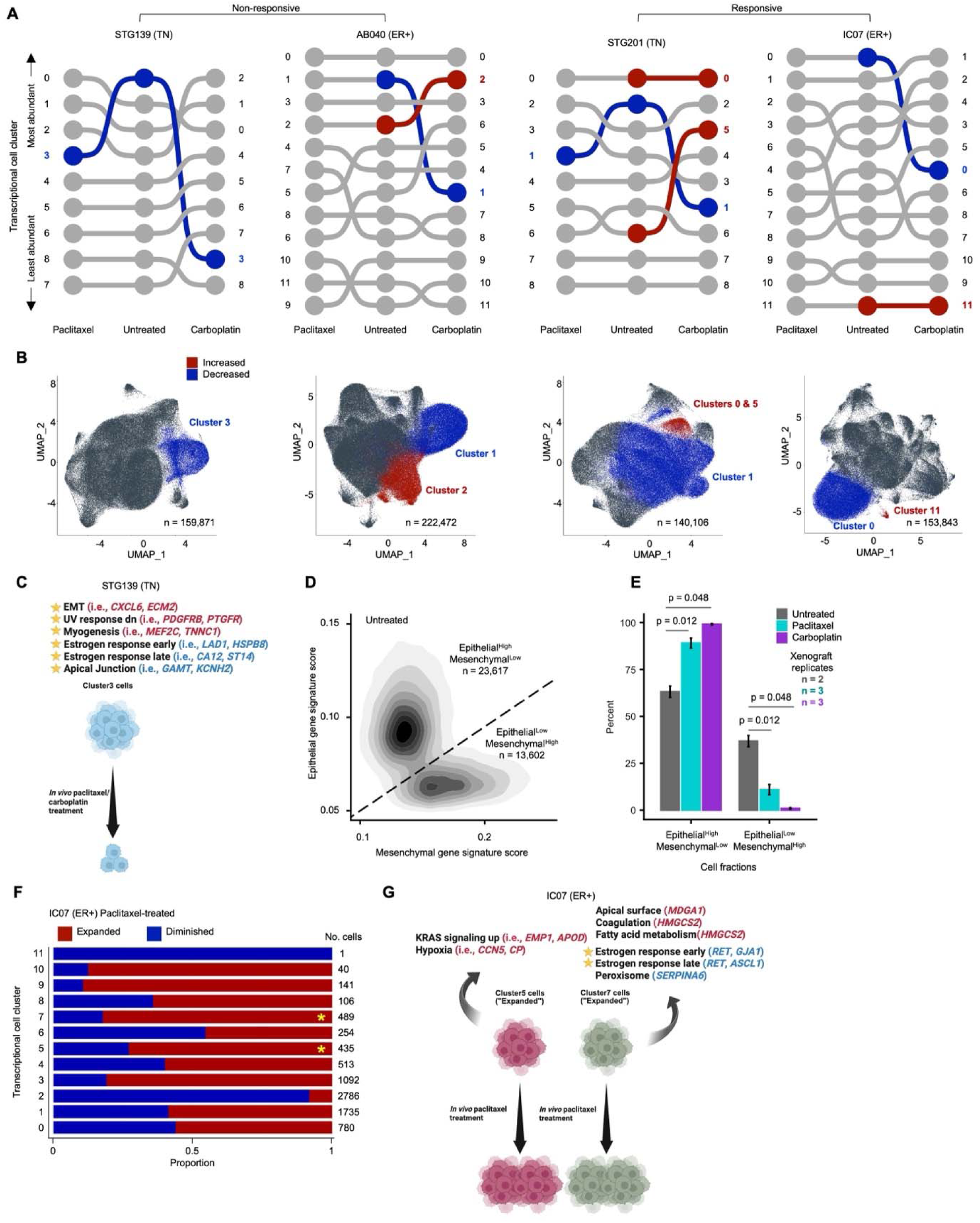
Transcriptional cell states exhibit differential sensitivity to chemotherapy. (**A**) Representation of cluster abundance across cohorts. Statistically significant increases and decreases in cluster proportion compared to untreated control are shown in red and blue, respectively, and correspond to Figure S7. (B) UMAP visualization of clusters that show a significant increase or decrease in proportion contribution with treatment, corresponding to (A). **(C)** Gene expression pathways that are enriched or reduced in cluster 3 for STG139 are indicated as red and blue, respectively. (D) Contour plot showing gene signature scores for individual cells from STG139 for untreated, paclitaxel and carboplatin treated cohorts. Diagonal line is drawn to separate cells into two fractions: epithelial-high/mesenchymal-low and epithelial-low/mesenchymal-high. **(E)** Percent of cells for each cohort from STG139 contributing to the two fractions. Error bars = standard error, and p-values from Welch^J^s t-test is shown. (F) Proportion contribution to each UMAP cluster by cells from expanded and diminished clones in ER+ model IC07. Significant difierences in proportion contribution to each cluster by Welch’s t-test are indicated with a yellow asterisk. Number of cells in each cluster listed on the far right. (G) Same as (C) but for significant cell clusters from (F) for IC07.

In the TN model STG139, cluster 3 was sharply depleted after both paclitaxel and carboplatin treatment (**Fig. 3A-B and S7A**). This cluster is characterized by elevated EMT signaling and suppression of estrogen response and apical junction pathways (**Fig. 3C**), consistent with a mesenchymal cell state that is preferentially eliminated by chemotherapy. Comparing untreated and treated xenografts from STG139 using epithelial and mesenchymal gene signature scores, we observed a clear shift toward cells with a high epithelial signature after treatment, with a corresponding depletion of cells with a high mesenchymal signature. Two phenotypically distinct populations emerged: Epithelial^High^/Mesenchymal^Low^ and Epithelial^Low^/Mesenchymal^High^ (**Fig. 3D**). Both paclitaxel and carboplatin significantly reduced the Epithelial^Low^/Mesenchymal^High^ fraction, with carboplatin producing the stronger effect (**Fig. 3E**). These results indicate that, in STG139, epithelial-like xenograft tumor cells are relatively drug tolerant, whereas mesenchymal-like xenograft tumor cells remain highly sensitive to both agents.

A contrasting pattern emerged in the second TN model, STG201. Here, EMT, hypoxia and angiogenesis pathways were enriched in cell cluster 5, a population that expanded significantly after carboplatin (but not paclitaxel) treatment (**Fig. 3A-B and S7-8**).

Additional context-dependent differences were observed in TN model STG201 and ER+ model AB040. TNF-alpha signaling was downregulated in cluster 1 of STG201, corresponding with a reduced contribution of treated cells across both therapies. In contrast, the same pathway was upregulated in cluster 2 of AB040, where it corresponded to an increased representation of cells following carboplatin (but not paclitaxel) treatment (**Fig. 3A-B and S7-8**).

Our lineage-resolved analysis enabled us to quantify how individual cell clones contributed to each transcriptionally defined cluster and to determine which clusters were more significantly associated with individual cell clones that diminished or expanded after chemotherapy. In the ER+ model IC07, clusters 5 and 7 showed a significantly higher contribution from expanded compared to diminished cell clones (**Fig. 3F**). Cluster 7 was marked by reduced enrichment for early and late estrogen-response pathways and increased expression of genes involved in cell polarity and fatty acid metabolism, while cluster 5 showed strong upregulation of KRAS signaling and hypoxia-related programs (**Fig. 3G**). Together, these findings suggest that carboplatin resistance in IC07 is associated with activation of KRAS-driven programs, metabolic rewiring, and hypoxia-adapted states rather than estrogen response signaling that is typically associated with ER+ breast cancer.

Taken together, these results show that chemotherapy does not act on tumors as a uniform pressure but instead drives selective expansion and loss of distinct transcriptional cell states. Responsive and intrinsically non-responsive clonal cell populations coexist within the same tumor, and their fate under treatment depends heavily on the cellular context in which key pathways are engaged. Notably, signaling programs such as EMT or TNF-α can promote sensitivity in one setting and resistance in another, underscoring that chemotherapy response is governed not by single pathways but by the transcriptional cell states in which those pathways operate. These findings highlight the complexity of cell clonal selection under treatment and point to context-specific cellular states as critical determinants of therapeutic outcome.

### Unbiased metaprogram discovery identifies stress-adaptive states linked to clonal resistance

To uncover gene programs associated with, or predictive of, cellular clonal responses to chemotherapy, we applied non-negative matrix factorization (NMF) to the scRNAseq datasets (**Methods**). Because NMF does not rely on predefined biological categories, it allows de novo identification of coordinated transcriptional modules. This analysis generated 12, 10, 15, and 12 metaprograms (MPs) in models STG139, STG201, AB040, and IC07, respectively (**Fig. 4A, S9**).

MP1 emerged as one of the most compelling metaprograms, identified in the ER+ model IC07 that responded to chemotherapy. MP1 expression was strongly enriched in expanded compared with diminished clones after carboplatin (p□<□2.2□×□10□^1^□; **Fig.□4B**). A similar but more modest pattern was seen in ER+ model AB040 (which was non-responsive to chemotherapy) (p□=□1□×□10□□), whereas no significant enrichment was observed in either TN model, suggesting that MP1 captures a stress-adaptive program characteristic of ER+ disease. MP1 comprises immediate-early response genes and regulators such as *DUSP1* and *KLF4*, which were consistently elevated in expression in expanded compared to diminished clones for both ER+ models. This pattern suggests rapid activation of stress-response circuitry and signaling re-equilibration that may confer platinum tolerance (**Fig.□4C**).

**Figure 4.**
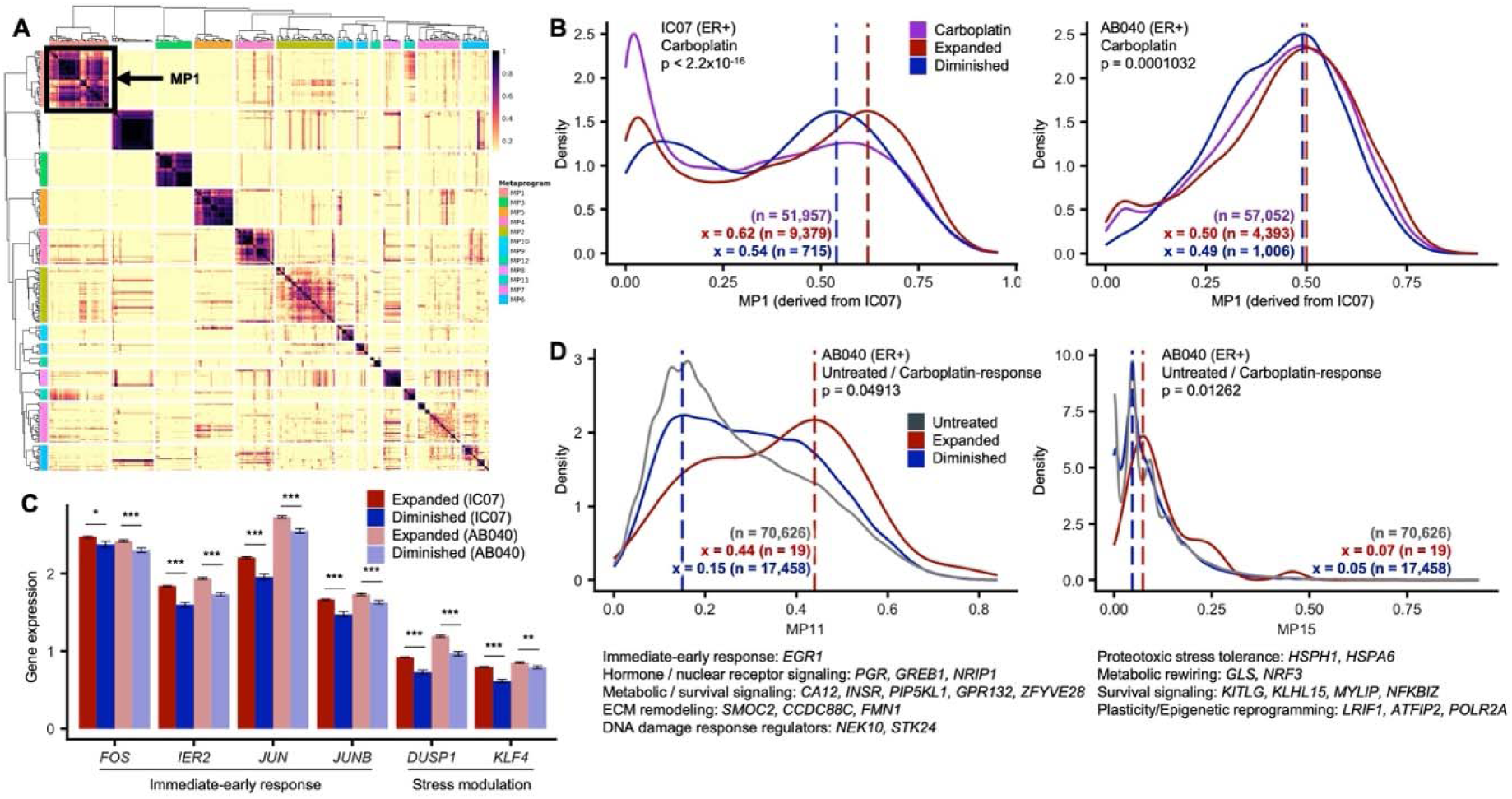
Metaprograms associated with response to Carboplatin. (**A**) Clustered heatmap of genes showing a positive linear correlation for ER十 model IC07, identifying 12 metaprograms. (B) Density plot ofMPl (derived from IC07 dataset) expanded and diminished clones after Carboplatin treatment (compared to the entire Carboplatin cohort, including barcoded and non-barcoded cells). P-values are from Welch’s t-test comparing expanded and diminished clones. X-axis values indicate the peak of the density distributions. (C) Barplot of individual genes showing significantly increased expression in expanded clones. Mean and standard error are represented with a black dot and error bars, respectively. Twotailed student’s t-test shown as * = p < 0.05, ** = p < 0.01, *** = p< 0.001. **(D)** Density plot ofMPl 1 and MP15 (derived from AB040 dataset) showing a statistically significant difference in expression in expanded and diminished clones identified in untreated AB040 xenografts (compared to all untreated cells, including barcoded and non-barcoded cells). Selected genes within each metaprogram are listed with their potential function.

We next asked whether MPs expressed at baseline could predict how individual cellular clones would respond to chemotherapy. Focusing on single-cell-derived clones detected across multiple xenografts, we examined clones with consistent behavior across treatments and labeled them in untreated xenografts according to their subsequent response to chemotherapy. This analysis revealed two additional metaprograms, MP11 and MP15 from ER+ model AB040, which were significantly enriched in clones that later expanded following carboplatin exposure (**Fig.□4D**). MP11 comprises genes involved in hormone and nuclear receptor signaling, metabolic and survival pathways, extracellular matrix remodeling, and DNA-damage response regulation. MP15 includes heat-shock proteins involved in proteotoxic stress tolerance, along with genes linked to metabolic rewiring, survival signaling, and epigenetic reprogramming. Collectively, these programs mark cell states with enhanced capacity to buffer oxidative and DNA-damage stress, while maintaining plasticity through metabolic and epigenetic adaptation.

Together, these findings demonstrate the power of integrating cell clonal lineage tracing with single-cell transcriptomics to pinpoint molecular programs that shape clonal fitness under chemotherapy. In ER+ models, *DUSP1* and *KLF4*-driven stress-response states emerge following carboplatin treatment and align with a slow-cycling, drug-tolerant persister-like phenotype. Moreover, the baseline expression of MP11 and MP15 identifies cellular clones predisposed to survive carboplatin, implicating metabolic, proteotoxic-stress, and epigenetic remodeling programs as key determinants of cell clonal resistance.

## DISCUSSION

We combine here clonal tracking analysis of single cell-derived clones with scRNAseq to gain new insights into cell clonal response to chemotherapy treatment. A key distinguishing feature of our study is that we track the growth and gene expression profiles of clones derived from single uniquely genetically barcoded cells, as opposed to genomic clones. Although single cell-derived clones and genomic clones may partially overlap, the molecular and clonal heterogeneity we identify in the former is driven predominantly by differences in transcriptional cell states. These cell states are shaped by multiple influences, including epigenetic regulation and the tumor microenvironment^16,17^.

A major finding reported herein is xenografts that grow at an exponential rate despite chemotherapy treatment, designated chemotherapy non-responsive, show the most dramatic changes in cellular clonal composition despite no perceptible impact on tumor growth. This is largely due to the emergence of previously minor cell clones that harbor a chemotherapy resistant phenotype to compensate for the decrease in size from dominant chemotherapy responsive cell clones. This constitutes direct evidence that responsive and non-responsive clonal cell populations can simultaneously co-exist within a tumor, and their proportional contribution to the tumor is dependent on the context. This is perhaps most evident in TN model STG139 where dichotomous phenotypic cellular populations can be identified based on epithelial and mesenchymal gene signatures. The Epithelial^High^/Mesenchymal^Low^ and Epithelial^Low^/Mesenchymal^High^ cell populations increase and decrease in prevalence following chemotherapy treatment, indicating they are differentially responsive. We and others have shown these two phenotypic cell populations can interconvert to maintain an equilibrium^18–21^ which we show here is perturbed by chemotherapy. This suggests that cell clone response to chemotherapy may be determined by the transcriptional and/or metabolic state of the cells derived from a single progenitor, which may exist as an equilibrium between chemotherapy responsive and non-responsive states. These findings have profound implications for how we approach cancer treatment because the molecular and cell clonal landscape of a tumor at the time of diagnosis may be completely altered after as little as two cycles of chemotherapy. This argues for the importance of repeat biopsies while on therapy to adjust treatment approach accordingly (i.e., adaptive therapy), as recently reported^22^.

Notably, we identify *DUSP1* and *KLF4* as putative markers of platinum chemotherapy resistance in luminal B/ER+ models of human breast cancer. *DUSP1* normally functions to dephosphorylate JNK and p38^23,24^, which are pro-apoptotic MAP kinases. The resulting effect is to blunt stress response by allowing the cells to survive oxidative damage and DNA-damage induced by platinum chemotherapy. *DUSP1* has previously been linked to platinum chemotherapy resistance^25,26^. As for *KLF4*, its function is complex and context-dependent, it often acts as a pro-survival transcription factor in cells treated with chemotherapy where it transcriptionally regulates genes involved in cell survival and repair, and induces a slow-cycling, stress-resistant phenotype consistent with a drug-tolerant persister state^27,28^. The I-SPY 1 neo-adjuvant therapy trial for patients with breast cancer who had tumor biopsies before treatment, after initiation of chemotherapy, and at time of surgery found significant changes in gene expression, including downregulation of proliferation and immune-related genes, consistent with a low-cycling, stress-modulatory state^29^, which is consistent with the stress modulatory effects of *DUSP1* and *KLF4*, and support the findings we report here.

We applied NMF to identify gene sets that can potentially be used to predict the response of individual cell clones to treatment. By annotating the cell clones in untreated xenografts based on the response of the same single cell-derived clones in treated xenografts, we identified 2 MPs that show increased expression in expanded compared to diminished clones with carboplatin treatment in ER+ breast cancer and implicate hormone and nuclear receptor signaling, metabolic rewiring and survival pathways, extracellular matrix remodeling and epigenetic reprogramming, consistent with a cellular state primed for plasticity, and that may facilitate molecular and metabolic adaptation to persist and survive under therapeutic stress.

The concept of a drug-tolerant persister state would be consistent with plasticity between chemotherapy responsive and non-responsive states, rather than a fixed resistance mechanism such as a genomic mutation. In models of lung cancer, resistant cellular clones were present before treatment but were rare, representing between 0.001 – 0.05% of the initial population of cells^30,31^, and a study by Oren et al using an expressed barcode library identified cycling and non-cycling cancer persister cells that originate from different pre-existing cell lineages^11^. These lineages express distinct transcriptional and metabolic programs even before drug treatment. These findings in lung cancer models are consistent with our finding that upon chemotherapy treatment, the dominant cell clones in the untreated condition can be quickly replaced by hundreds of emerging cell clones that harbor a chemotherapy resistance phenotype. The magnitude of change to the cellular clonal landscape varied between PDTX models, consistent with the heterogeneity of response and clonal composition expected between different breast cancer subtypes. However, the transcriptional and metabolic programs associated with the cell clones harbouring a chemotherapy resistance phenotype were also pre-existing and expanded upon treatment.

By coupling clonal lineage tracing with single-cell expression profiling, we uncover a level of cellular clonal complexity and cell state plasticity that cannot be appreciated from bulk analyses. A central implication is that resistance frequently originates from rare, pre-existing cell clones that evade detection with conventional sequencing. This limits the ability of bulk tumor profiling to guide therapy selection or anticipate resistance. Approaches incorporating serial and on-treatment biopsies, together with PDTX models deployed as patient surrogates^32^, may be required to capture emerging resistant populations and better inform treatment decisions.

## RESOURCE AVAILABILITY

### Lead contact

Requests for further information and resources should be directed to and will be fulfilled by the lead contact, Long V. Nguyen.

### Materials availability

This study did not generate new PDTX models. All unique reagents generated in this study are available from the lead contact with a completed materials transfer agreement.

### Data and code availability

Processed count matrices from scRNAseq can be accessed through Zenodo (<u>10.5281/zenodo.17818897</u>). Reviewers can access the restricted files anonymously using the following link: https://zenodo.org/records/17818898?token=eyJhbGciOiJIUzUxMiJ9.eyJpZCI6IjVjMTdiN2ViLWM3N2ItNDRkMy04NGJmLTVhNGMyZjc2YTYwNCIsImRhdGEiOnt9LCJyYW5kb20iOiIzZDEyYzAyNjNlNDNiM2Q5MjQ5Yzc4M2U5MjUyMjZhZCJ9.Jou5AZp7qNQVJNMOoLJWl0crvs2oOdwSQ0vLuvOuAQ3Msu2gBbBBw_N7s6BjWTFm0fwjeFRLdtNa_gT0KwAIhA. These files will be published and unrestricted upon publication. Code to reproduce the computational analysis in this study are available at https://github.com/lvnguyen-lab/Chemo-response-expressed-barcoding and will be made available to reviewers upon manuscript submission.

## ACKNOWLEDGEMENTS

S.K. was supported by the Canadian Cancer Society Research Training Award (708339). J.W. was supported by the University Health Network Canada Leads Program. L.V.N. was supported by the Princess Margaret Cancer Foundation, a Temerty Faculty of Medicine Hold’em for Life Early Career Professor in Cancer Research Award, the Canadian Institutes of Health Research (PJT-204015 and PJF-204191), a MOHCCN Clinician-Scientist Award (3255-02), an ESMO Translational Research Fellowship and an ASCO Conquer Cancer Young Investigator Award. D.G.-R. is funded by the Cambridge Commonwealth, European and International Trust. O.M.R. was supported by the UKRI grant MC_UU_0002/16. F.G. was supported by Princess Margaret Cancer Foundation, Ontario Institute for Cancer Research Investigator Award, the Canadian Institutes of Health Research, and the Natural Sciences and Engineering Research Council of Canada. C.C. was supported by funding from CRUK (grant numbers A17197, A27657 and A29580), an NIHR Senior Investigator Award (grant number NF-SI-0515-10090) and a European Research Council Advanced Award (grant number 694620). We are grateful for the generosity of all the patients who donated samples for the development of the tumor xenograft models; and the CRUK Cambridge Institute Core Facilities (Genomics, Flow Cytometry, Histopathology and Biorepository) for support during the execution of this project. Figures were created using Biorender.com.

## AUTHOR CONTRIBUTIONS

L.V.N. and C.C. conceived the study and wrote the manuscript together with S.K. Data analysis was led by S.K. and L.V.N. with input from J.W., J.Y.C., D.G.-R., O.M.R., and F.G. Tumor xenograft experiments were designed and led by L.V.N. with input from S.-F.C., G.L., and A.J.W.L. All authors read and approved the manuscript.

## METHODS

### PDTX barcoding

A single cell suspension of PDTX cells were genetically labeled with a lentiviral barcode library, based on the pLARRY barcoding system^12,33^, such that each transduced cell contains a unique and permanently integrated DNA-based barcode sequence that is also expressed into mRNA transcripts and thus compatible with single cell RNA sequencing (scRNAseq)^21^. Two basal subtype triple-negative breast cancer PDTX models STG139 and STG201, and two luminal B subtype ER+/HER2-models AB040 and IC07 were used^34^. PDTX fragments were dissociated into a single cell suspension (Miltenyi Biotec, Cat. no. 130-095-929) and depleted of mouse cells (Miltenyi Biotec, Cat. no. 130-104-694). Tumor cells were then transduced with lentiviral barcode library BC1 as previously described^21^. We targeted a transduction efficiency of ≤30% to reduce the likelihood of multiple integrations. A small fraction of transduced cells were maintained *in vitro* for 48-72hrs for flow cytometry analysis to estimate transduction efficiency, as reported in **Table S5**. Out of 1070 and 1091 unique barcodes detected in models STG201 and AB040, 1 (0.0009%), and 13 (0.01%) sets of multiple integrations were detected. No multiple integrations were detected for STG139 and IC07. This indicates the frequency of multiple integrations based on our lentiviral transduction conditions was ≤ 0.01% (**Table S6**). Only one barcode within a set of multiply integrated barcodes was retained in the final dataset, and this was corrected in our DNA amplicon sequencing dataset as well. Transduced cells were immediately used for mouse engraftment without a period of *in vitro* culture to minimize the likelihood of cell division, to ensure all clones detected *in vivo* were derived from single uniquely barcode-labeled cells.

### Mouse experiments and chemotherapy treatment

Transduced cells were resuspended in 100ul of 50% Matrigel and 50% RPMI growth media and injected subcutaneously into 8–12-week-old female NSG mice. Weekly tumor measurements were made using calipers and tumors harvested upon reaching 1500 mm^3^ or another humane endpoint. Time to tumor harvest ranged from 64 to 127 days and 84 to 180 days for primary and secondary xenografts, respectively (**Table S5**). Tumors from primary xenografts were minced into small pieces < 3 mm^3^ and viably frozen in FBS with 10% DMSO. To established secondary xenografts, frozen vials of PDTX fragments were rapidly dissociated and immediately injected subcutaneously into NSG mice as previously described^21^. Nine secondary xenografts were established from 1 primary xenograft for each PDTX model (n = 3 for each cohort, untreated, paclitaxel-treated and carboplatin-treated). For secondary xenografts only, once the xenografts for a given PDTX model reached an average size of 100 mm^3^, mice were assigned to study cohorts and treatment initiated. To reduce bias and ensure balance between study cohorts, the xenografts were ordered by tumor volume, and the spiral method used to assign mice to cohorts. Carboplatin (Selleckchem, Cat. no. S1215) was administered at a dose of 40mg/kg intravenously every 3 weeks for 2 doses. Paclitaxel (Selleckchem, Cat. no. S1150) was administered at a dose of 7 mg/kg intravenously weekly for 6 doses. Both treatments were designed to mimic 2 cycles of treatment as would be given in patients. Following chemotherapy treatment, xenografts were harvested upon reaching 1500 mm^3^ or another humane endpoint. Time from treatment completion to tumor harvest is indicated in **Fig. 1B**. Specially, tumors for PDTX models IC07 (both paclitaxel and carboplatin treated cohorts) and AB040 (2 of 3 xenografts from the paclitaxel treated cohort) were harvested more than 30 days after treatment completion. One untreated xenograft from STG139 was not included in the dataset as sequencing had failed.

### Barcode DNA amplicon sequencing and analysis

Xenografts were harvested for DNA amplicon sequencing and scRNAseq analysis when tumor size reached 1.5 mm^3^ (**Table S5**). Barcode DNA amplicon sequencing and data processing were performed as previously described^21,35^. Sequencing was performed on the Illumina MiSeq for 75bp paired-end sequencing. On average each sequencing run yielded 2 x 10^7^ paired-end reads, resulting in between 5 x 10^5^ to 1 x 10^6^ paired-end reads per sample analyzed.

### Classification of response to treatment

Models were assigned a bulk response to treatment using custom code provided: “tumor_volume_bulk_response.ipynb”. This script uses tumor volume measurements over time for all three secondary cohorts and calculates the growth rate of each xenograft in the last week of treatment (or the last week prior to harvest if the PDTX was harvested during the treatment window). The script then performs a one-tailed t-test to determine whether the treated growth rate is significantly below the untreated growth rate. Models for which the treated growth rates in the last week of treatment were significantly below the untreated growth rates were classified as responsive, while models for which this was not the case were classified as non-responsive.

Clones in treated xenografts were assigned a response to treatment using custom code provided: “amplicon_analysis.ipynb”. This script reads the processed amplicon files, compares the barcode clones present within each PDTX model to determine how they change in size between treated and untreated xenografts. Clones that decrease in size by at least 30% compared to the average clone size in untreated xenografts are classified as “diminished”. Clones that increase in size by at least 20% compared to the average clone size in untreated xenografts are classified as “expanded”. Clones that are neither “diminished” nor “expanded” are classified as “minimal change”. We also classified clones in untreated xenografts based on how the same clone detected in treated xenografts responded to paclitaxel or carboplatin treatment. Clones in untreated xenografts were classified as “discrepant” if the same clone was classified as “expanded” and “diminished” in different treated xenograft replicates.

### scRNAseq and data processing

Samples were prepared and sequenced as previously described^21,35^. 10X Chromium scRNAseq was performed using standard 3’ v3.1 chemistry for primary xenografts for PDTX models STG201 and IC07 to target recovery of 1 x 10^4^ cells. All other primary and secondary xenografts were analyzed using the 3’ v3.1 HT chemistry to target recovery of 2 x 10^4^ cells. Sequencing was performed on the Illumina NovaSeq S4 flowcell targeting 2 x 10^4^ reads per cell. For data processing, demultiplexed FASTQ files were processed by *cellranger* (v8.0.1)^36^ count command and Seurat (v5)^37^ objects were created from each sample. A QC threshold was applied to filter cells based on the percent mouse UMIs, number of UMIs, number of features, and percent mitochondrial UMIs; and scDblFinder^38^ was used to give each cell a doublet score and identify any clusters composed primarily of presumed doublets (**Table S7**).

Barcodes were associated with single cell RNA profiles as previously described in our protocol^35^. All barcode clones were assigned a response to treatment based on DNA amplicon sequencing analysis using custom code provided: “parse_barcodes_for_scRNAseq.ipynb”. This script reads the barcode association csv files from the process described in our protocol^35^ and the barcode response output from the “amplicon_analysis.ipynb” Jupyter notebook. Cases where multiple barcodes were associated with a single cell were classified as multiplets or multiple integrations. Sets of barcodes assigned to the same cell were classified as multiple integrations if they were detected together in at least 3 separate cells within a PDTX model, at least 10% of the cells containing any of the barcodes have all barcodes in the set, and if the response of all barcodes in the set to treatment was consistent. If a set of barcodes did not meet these criteria, it was classified as a multiplet. Multiplets were excluded from response-based analyses (**Table S7**). For multiple integrations, the barcode with the most sequencing reads identified from the “amplicon_analysis.ipynb” output was assigned as the final barcode for all cells with that set of multiple integrations (**Tables S6 and Fig. S10**). Cells where multiple barcode integrations could not be clearly defined as a set of multi-integrated barcodes were excluded from clonal analysis. This information was appended to the metadata in each Seurat object containing scRNAseq data.

Samples from each model were combined into a single Seurat object per model, and each model was transformed using Seurat’s SCTransform function and clustered based on gene expression patterns. Barcodes were not taken into account for clustering, and the impact of genes involved in hypoxia, IFN response, ncRNA, and stress response (**Table S8**) as well as cell cycle, mouse genes, and mitochondrial genes was regressed out prior to clustering. Clusters were optimized using Clustree (v0.5.1)^39^ to ensure cluster stability (**Fig. S4B**). Resolutions from 0.1 to 0.6 and k-values from 15 to 30 were evaluated, and optimal values for each model were chosen (**Tables S9 and S10**).

### ScRNAseq differential gene and pathway expression analysis

ScRNAseq clusters were analyzed to determine statistically significant differences in their composition with regards to treated and untreated cells and response to treatment. Each treatment was considered separately when compared to the untreated secondary cohort, and response to treatment was considered only within each secondary cohort. The R package *ggplot2*^40^ was used to visualize cluster composition via bar graph, and statistical analysis was performed using replicates to determine significant compositional differences between clusters.

Differential gene expression was determined between combinations of Seurat clusters, secondary cohorts, and the combination of secondary cohort and response to treatment using Seurat’s FindMarkers function with the MAST method. Parameters logfc.threshold, min.pct, and min.diff.pct were set to –Inf to obtain the full gene list regardless of log fold change and a rank for each gene was calculated by multiplying log2 fold change by –log10(adjusted p-value) and differentially expressed genes were visualized using *EnhancedVolcano*^41^. Gene set enrichment analysis (GSEA) was then performed using the resulting ranked list of genes against biological process gene ontology gene sets^42,43^ with the *ClusterProfiler*^44^ R package. The R package *GOSemSim*^45^ was used to determine semantic similarity between resulting GO pathways, and GSEA results were visualized using the R package *enrichplot*^46^. The R package *simplifyEnrichment*^47^ was used to cluster GSEA results, and clusters were given names by manual inspection of the pathways included in each cluster.

### Non-negative matrix factorization (NMF) analysis

NMF was performed on each model using the R package *GeneNMF*^48^ on the RNA assay with k=4:9, minimum average log-expression value of 0.05, cosine metric, 0.8 weight explained, min confidence 0.7, specificity weight 8, and a number of metaprograms (MPs) that resulted in significant differences between populations. NMF resulted in 12, 10, 15, and 12 MPs for models STG139, STG201, AB040 and IC07, respectively. MPs were maintained for analysis if they contained at least 10 genes, a Silhouette score of at least 0.3, and if they distinguished interesting sample populations based on response to treatment or cohort (**Table S11**). The R package *UCell*^49^ was used to score each dataset on the useful MPs.

## SUPPLEMENTARY FIGURES

**Figure SI.**
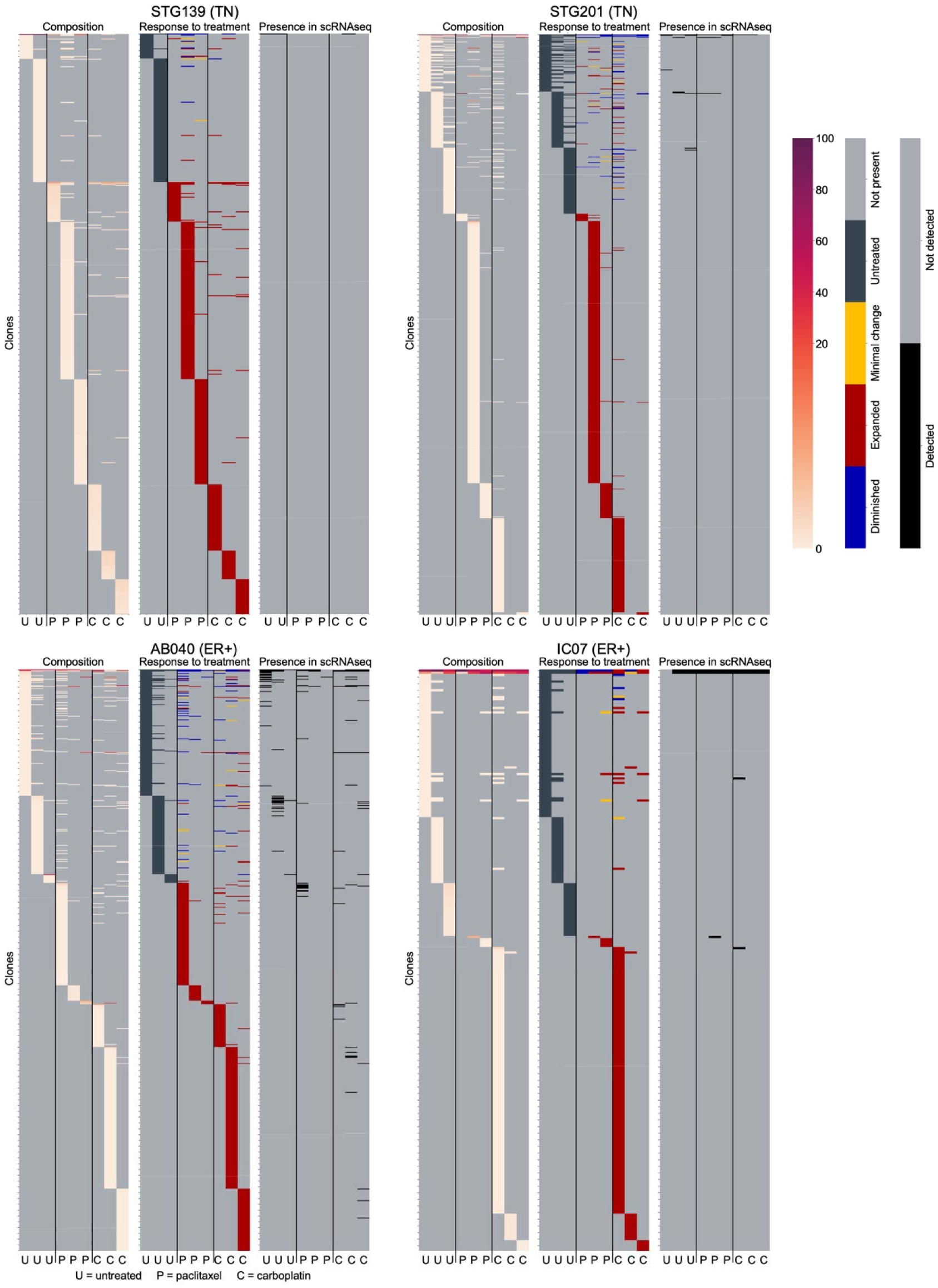
Overview of all clones detected by DNA amplicon sequencing and scRNAseq. Heatmaps with clones ordered top to bottom by most to least prevalent, and xenograft replicates ordered by column. Heatmaps show each individual clone in the dataset based on the % contribution to all barcoded cells in the xenograft (left), response to chemotherapy (middle), and whether it was detected by scRNAseq (right).

**Figure S2.**
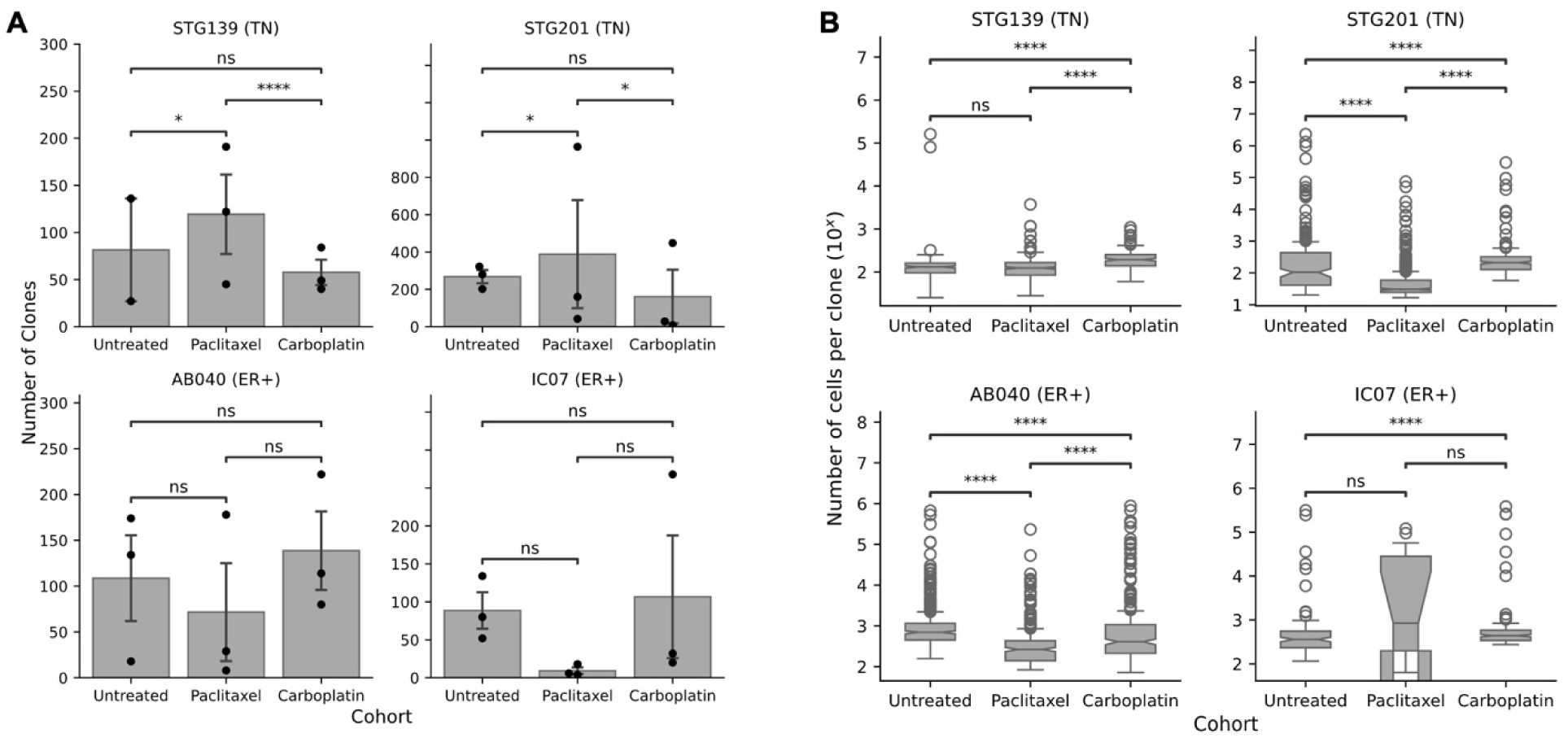
Number and size of clones across treatment cohorts. (**A**) Number of clones per xenograft. Points represent individual xenografts, with bar and error bars representing the mean and standard error respectively; n=2 PDTX for STG139 untreated and n=3 PDTX for all other conditions. (B) Clone size distribution across cohorts and models. For models STG139, STG201, AB040, and IC07 untreated, paclitaxel-treated, and carboplatin-treated respectively n=163, 3 58, 173, 402, 583, 242, 326, 215, 416, 133, 14, and 160 clones comprising 3 biological replicates. For **(A)** and **(B)** Two-tailed student^’^s t-test shown as ns = not significant, * = p < 0.05, ** = p < 0.01, *** = p< 0.001, **** =p< 0.0001.

**Figure S3.**
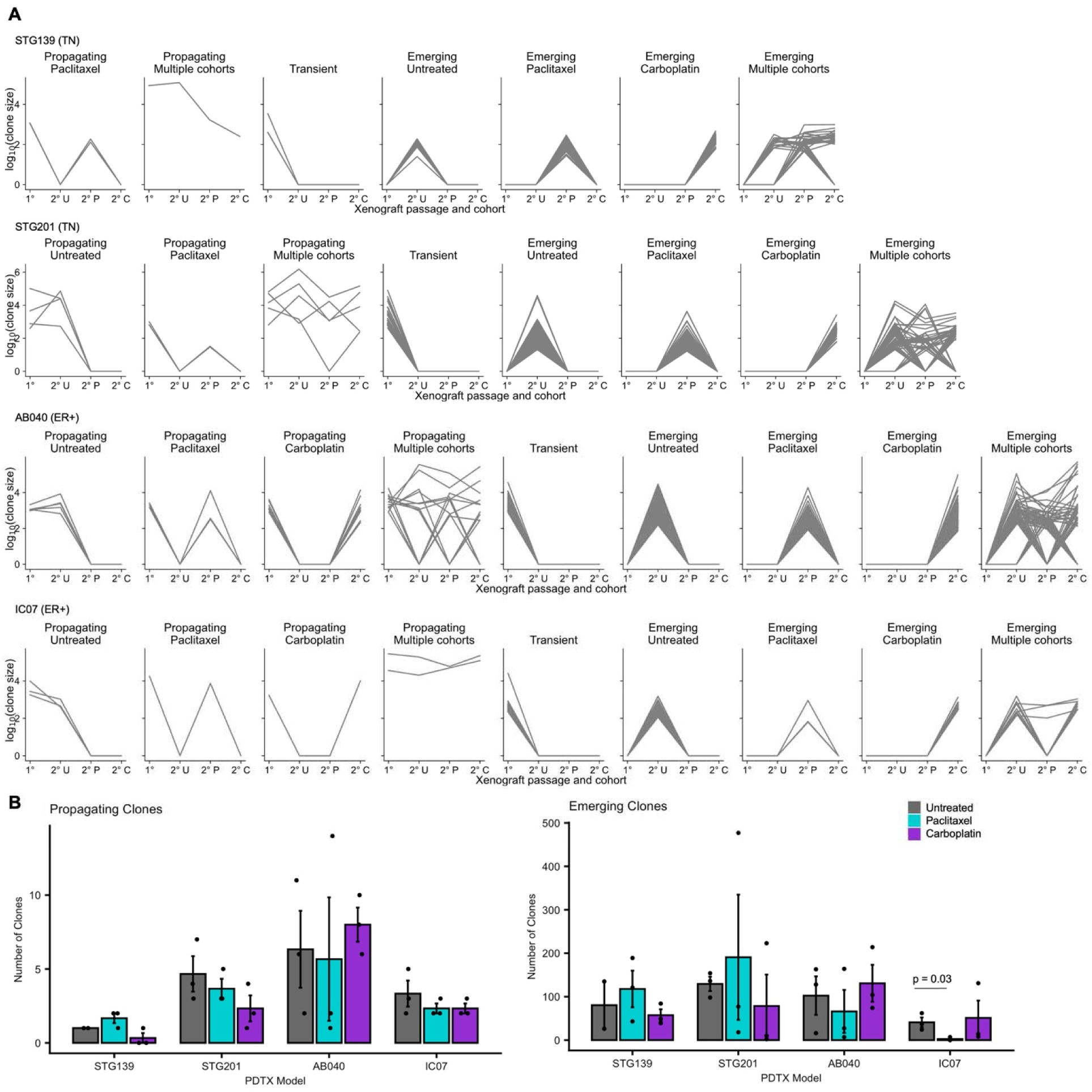
Clonal growth patterns detected by DNA amplicon sequence across treatment cohorts. (**A**) Each clone is represented by a single line indicating its size across treatment cohorts. The clones are labeled as propagating, transient or emerging. (B) Number of propagating (left) and emerging clones (right) across treatment cohorts. Only statistically significant differences compared to untreated (control) are shown. Student’s two-tailed t-test. Error bars = standard error.

**Figure S4.**
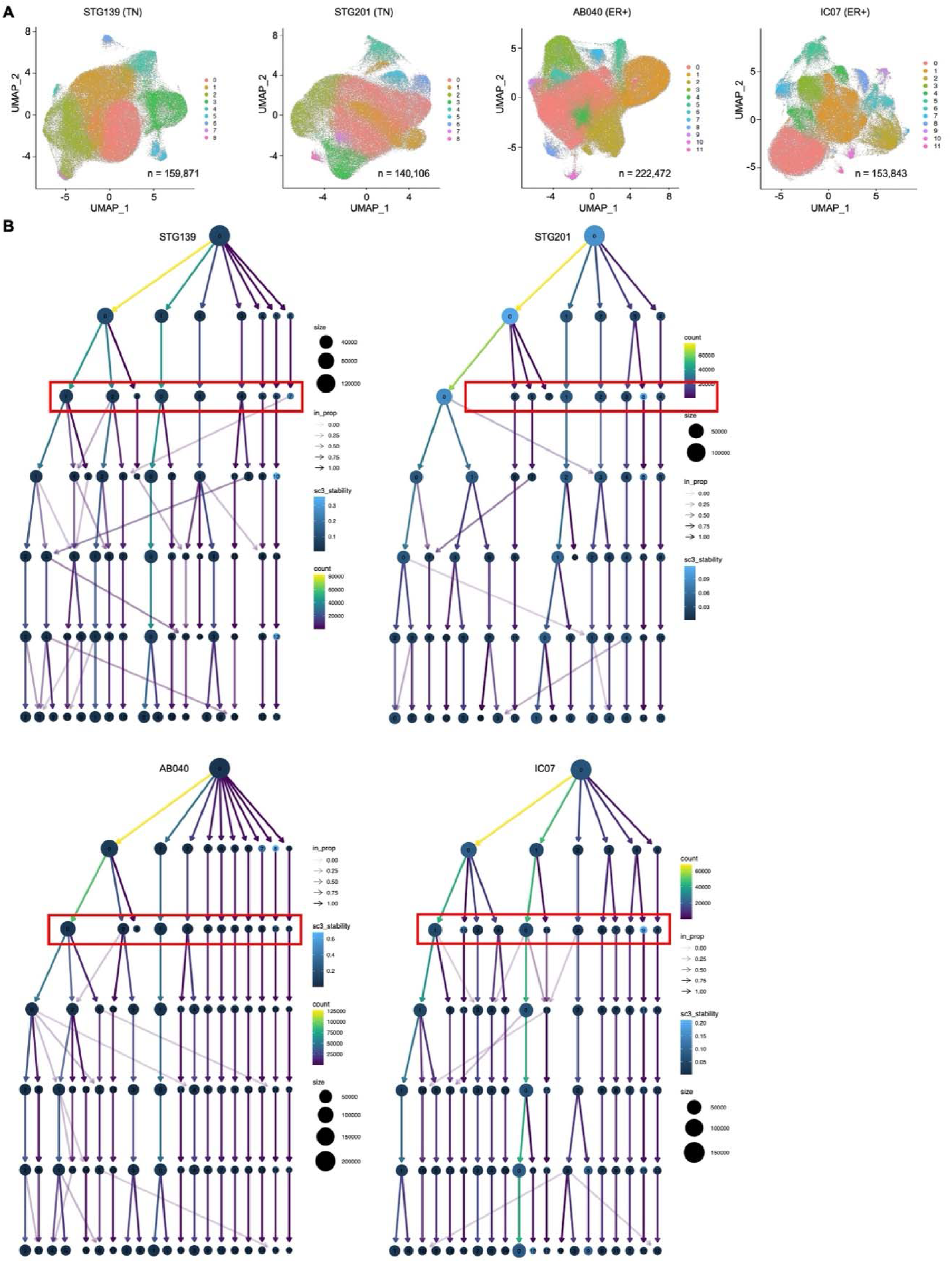
Clustering of single cell RNA profiles from scRNAseq. (**A**) UMAP of each PDTX model, colored by cluster **(B)** Clustree diagrams showing cluster assignments from Seurat clustering of our scRNAseq dataset at k = 20, 15, 25, and 15 respectively for models STG139, STG201, AB040, and IC07 at resolutions ranging from 0 to 0.6 in increments of 0.1. Red boxes indicate the ideal number of clusters beyond which there is instability of cluster assignments, indicating over clustering of the dataset.

**Figure S5.**
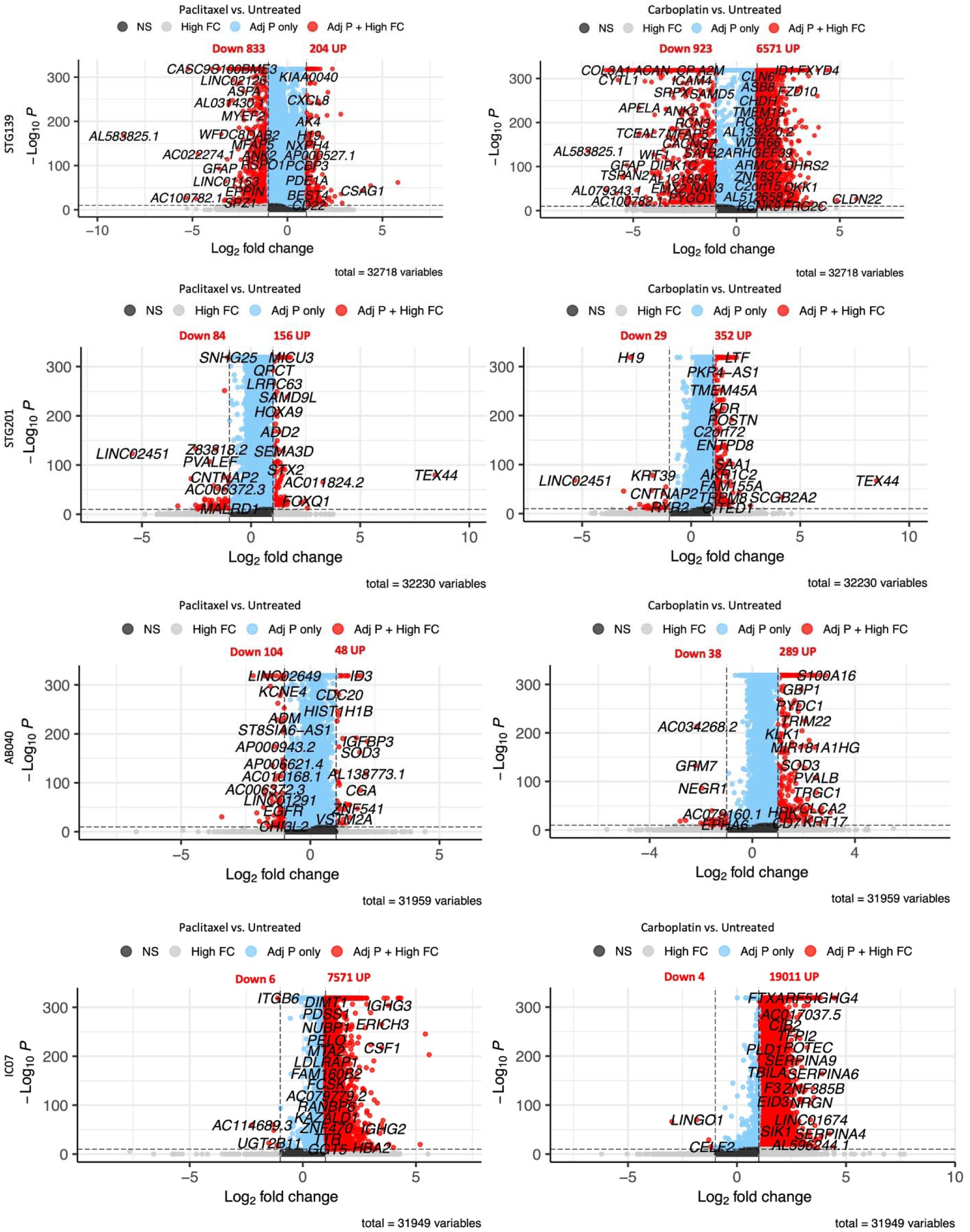
Volcano plots from differential expression analysis comparing paclitaxel or carboplatin treated xenografts to untreated xenografts. Genes that show a statistically significant difference in expression are shown in red, and those that do not are shown in gray (below 10 “ ° adjusted p-value for significance), blue (less than l-log2fbld change).

**Figure S6.**
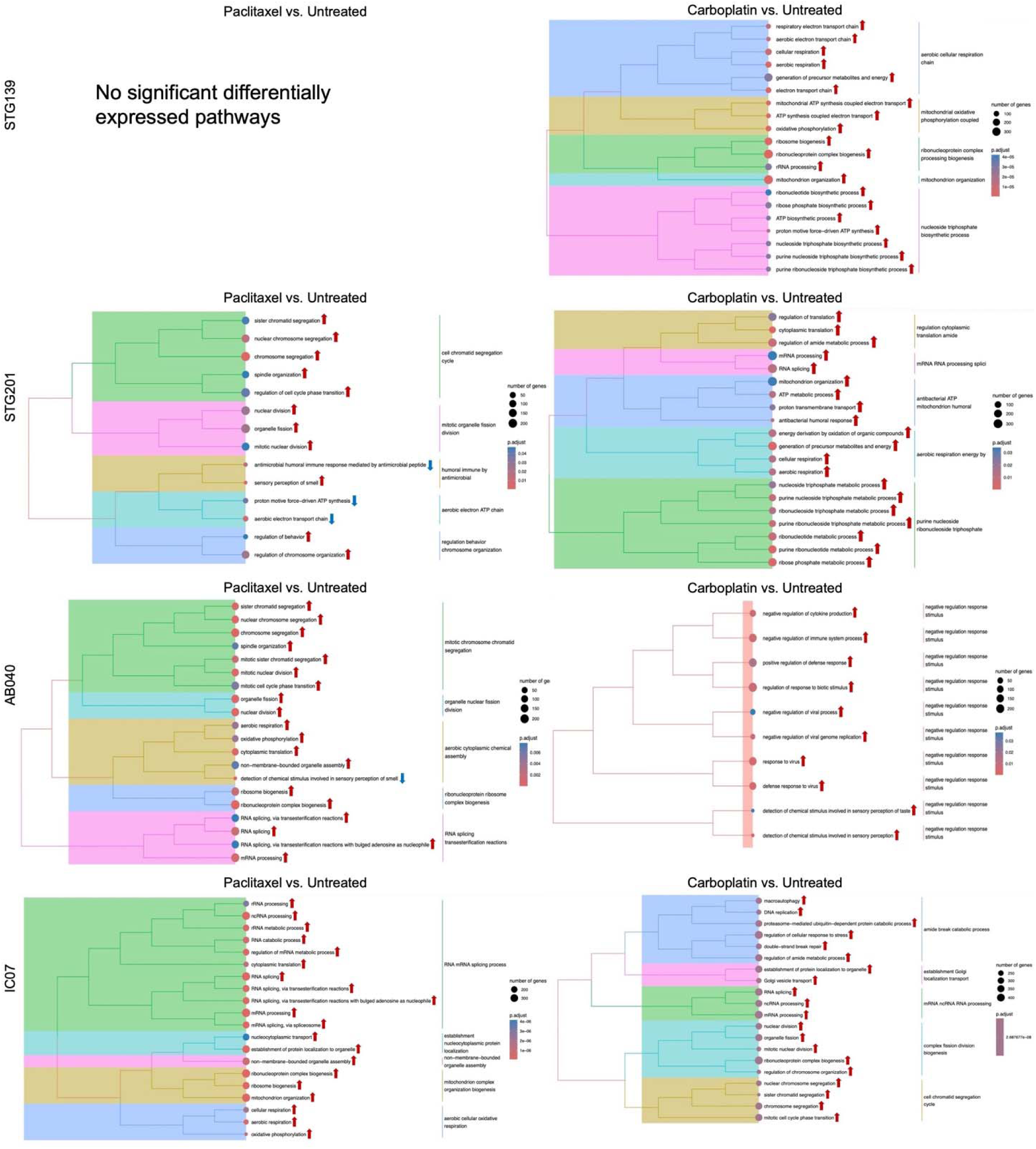
Top 20 significantly differentially expressed pathways comparing paclitaxel or carboplatin treated xenografts to untreated xenografts. All pathways shown are significant in their differential expression, with adjusted p-value represented by the colour and number of genes in the pathway by the size of the dots. Arrows indicate whether the pathway was up– or down-regulated compared to untreated control. Pathways are further grouped into similar processes indicated on the right. Results are based on differentially expressed genes from Fig. S5.

**Supplementary Figure S7.**
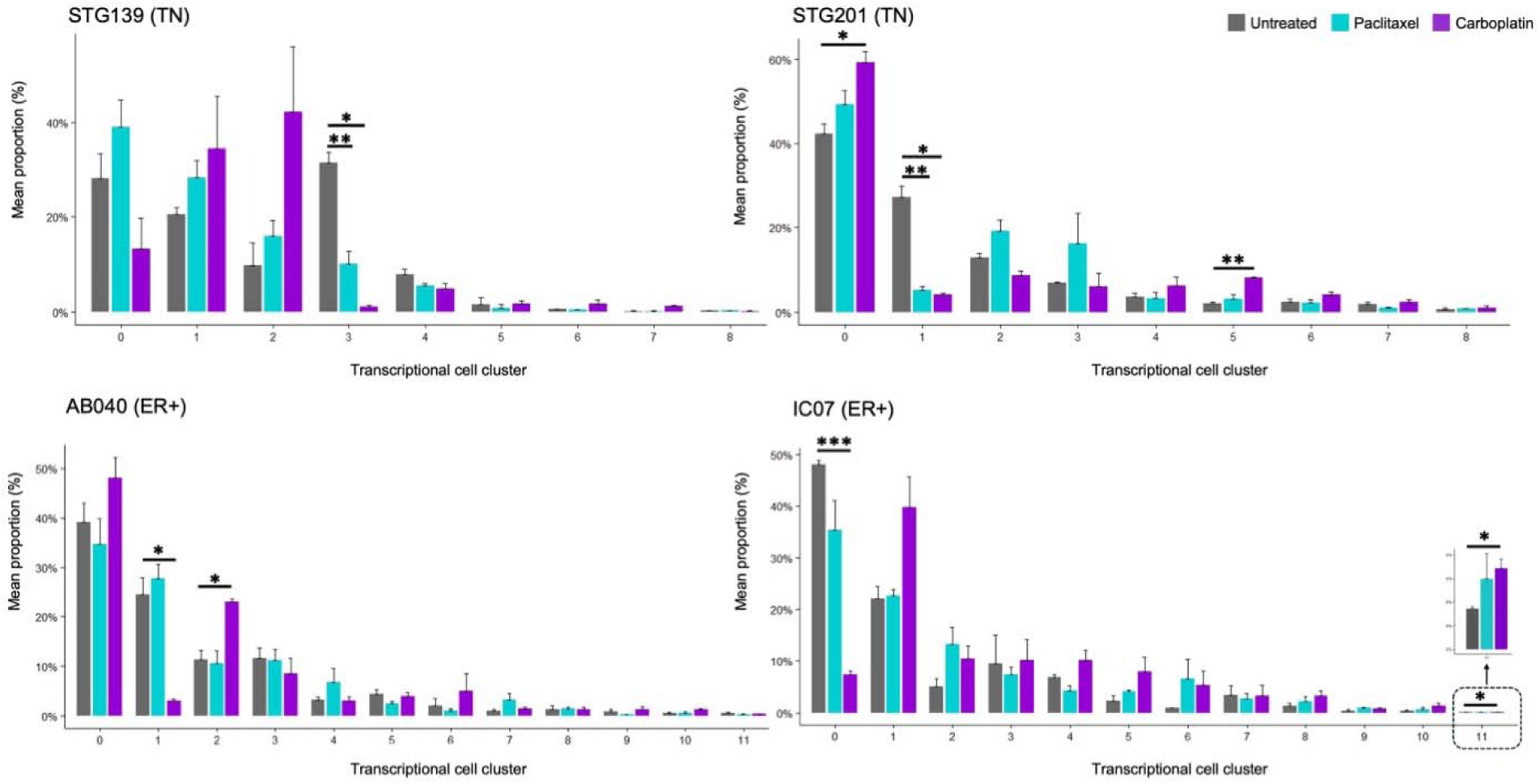
Mean proportion of cells across transcriptional cell clusters by cohort. Calculated for each xenograft replicate, as the proportion of cells in one cluster out of all cells for that xenograft. Error bars = standard error and p-values are from Welch’s t-test. * = p < 0.05, ** = p< 0.01, *** = p < 0.001.

**Figure S8.**
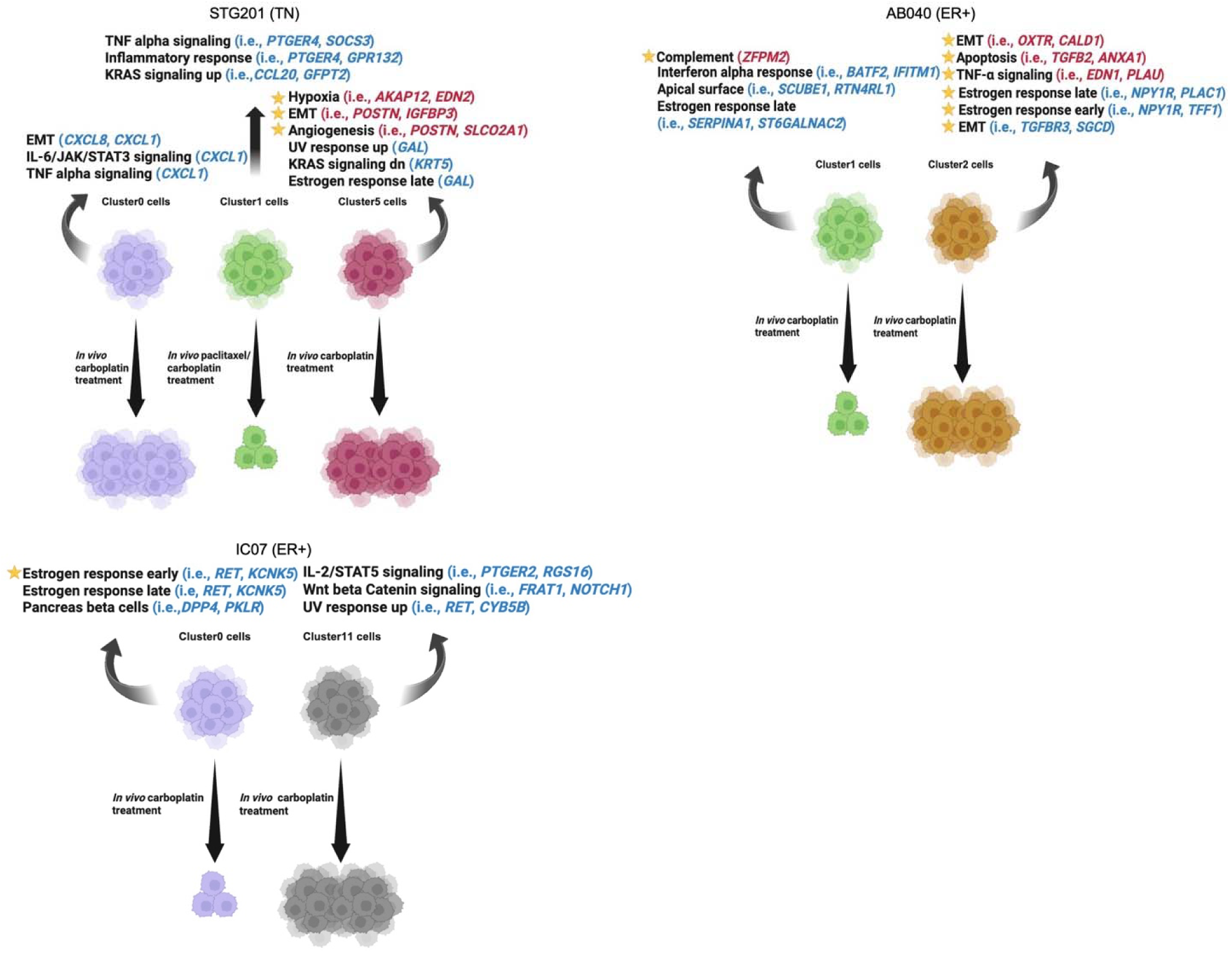
Gene expression pathways defining cell clusters of interest. Genes corresponding to cell clusters with statistically significant increased and decreased expression are indicated as red and blue, respectively. Enriched pathways are indicated with a yellow star symbol.

**Figure S9.**
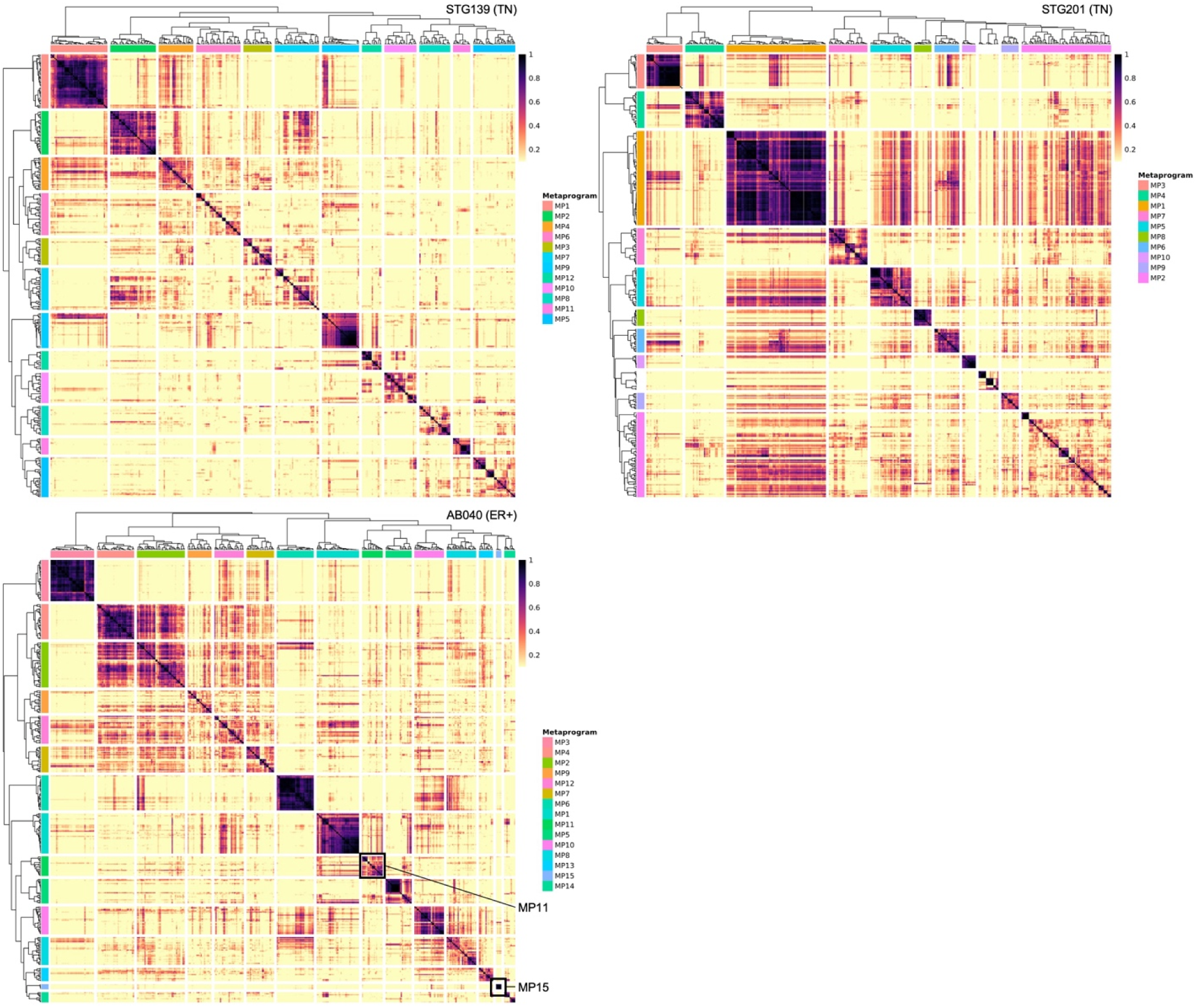
Clustered heatmaps from non-matrix matrix factorization (NMF) analysis. NMF analysis was performed on the scRNAseq dataset separately for each PDTX model. Defined metaprograms are shown, corresponding to gene sets that show a consistently strong correlation.

**Figure S10.**
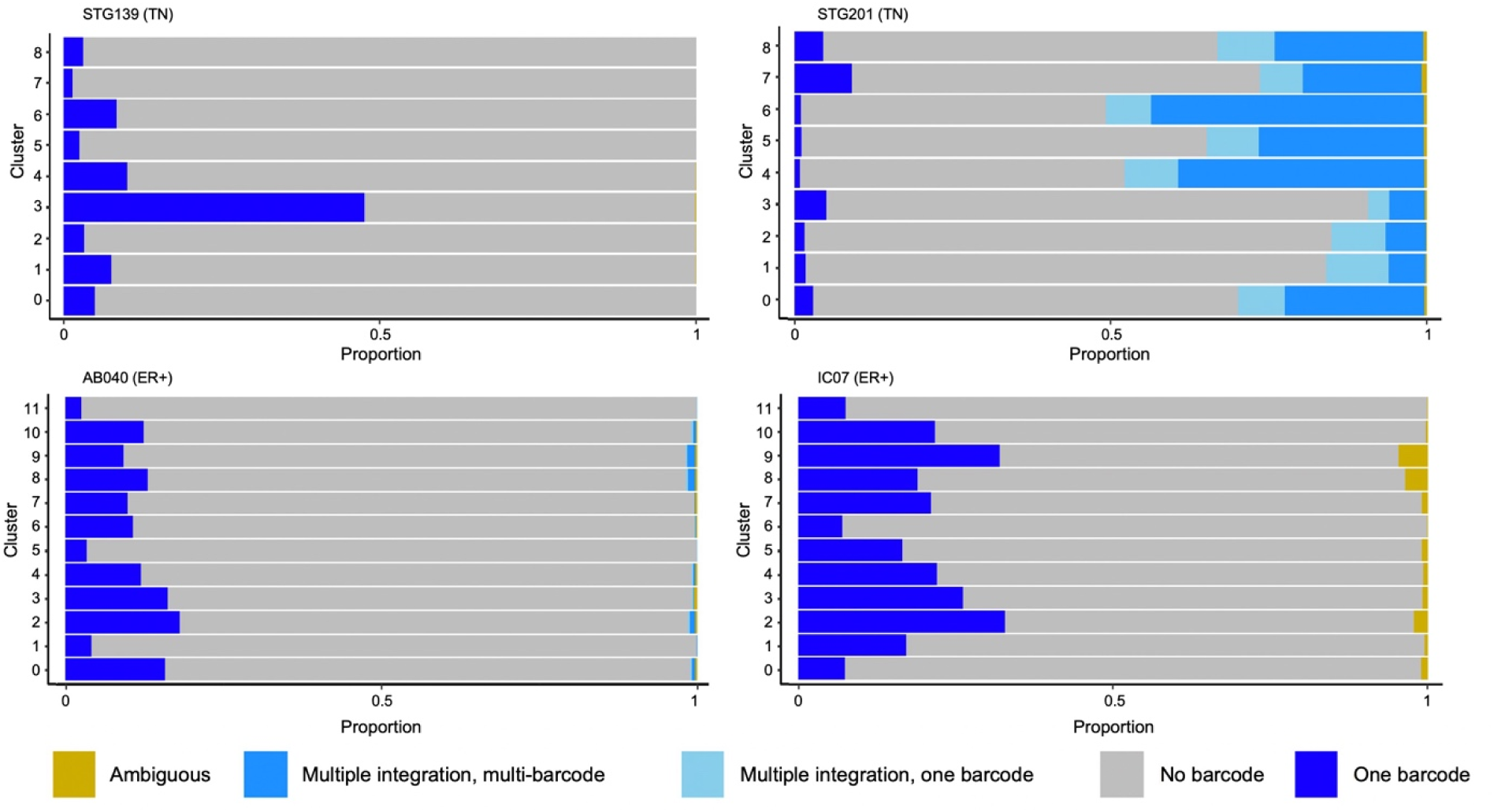
Proportion of scRNAseq dataset with barcodes detected. Barplots shown represent the distribution of barcoded cells across cell clusters. The proportion of cells with multiple barcode integrations is also shown, based on whether only one barcode of the set of multi-integrated barcodes or multiple barcodes of the set of multi-integrated barcodes were detected in the cell.

## SUPPLEMENTARY TABLES

(See excel files – except for Table S3 below)

**Table S3.**
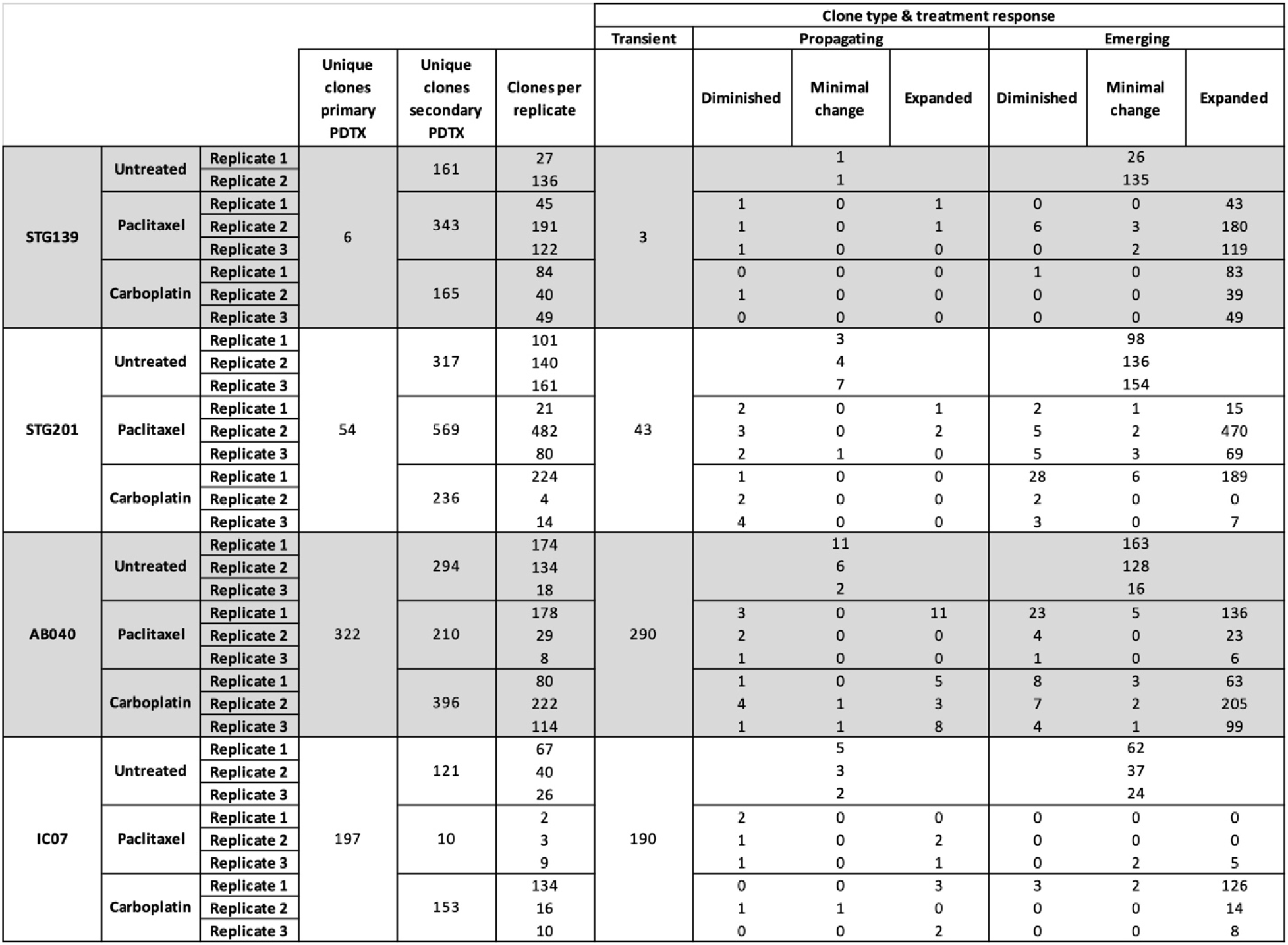
Overview of all clones detected by DNA amplicon sequencing.

## REFERENCES

1. Aalam, S.M.M., Nguyen, L.V., Ritting, M.L., and Kannan, N. (2024). Clonal tracking in cancer and metastasis. Cancer Metastasis Rev. 43, 639–656. 10.1007/s10555-023-10149-4.

2. Nguyen, L.V., Cox, C.L., Eirew, P., Knapp, D.J.H.F., Pellacani, D., Kannan, N., Carles, A., Moksa, M., Balani, S., Shah, S., et al. (2014). DNA barcoding reveals diverse growth kinetics of human breast tumour subclones in serially passaged xenografts. Nat. Commun. 5, 5871. 10.1038/ncomms6871.

3. Nguyen, L.V., Makarem, M., Carles, A., Moksa, M., Kannan, N., Pandoh, P., Eirew, P., Osako, T., Kardel, M., Cheung, A.M.S., et al. (2014). Clonal analysis via barcoding reveals diverse growth and differentiation of transplanted mouse and human mammary stem cells. Cell Stem Cell 14, 253–263. 10.1016/j.stem.2013.12.011.

4. Schepers, K., Swart, E., van Heijst, J.W.J., Gerlach, C., Castrucci, M., Sie, D., Heimerikx, M., Velds, A., Kerkhoven, R.M., Arens, R., et al. (2008). Dissecting T cell lineage relationships by cellular barcoding. J. Exp. Med. 205, 2309–2318. 10.1084/jem.20072462.

5. Serrano, A., Berthelet, J., Naik, S.H., and Merino, D. (2022). Mastering the use of cellular barcoding to explore cancer heterogeneity. Nat. Rev. Cancer 22, 609–624. 10.1038/s41568-022-00500-2.

6. Merino, D., Weber, T.S., Serrano, A., Vaillant, F., Liu, K., Pal, B., Di Stefano, L., Schreuder, J., Lin, D., Chen, Y., et al. (2019). Barcoding reveals complex clonal behavior in patient-derived xenografts of metastatic triple negative breast cancer. Nat. Commun. 10, 766. 10.1038/s41467-019-08595-2.

7. Lu, R., Neff, N.F., Quake, S.R., and Weissman, I.L. (2011). Tracking single hematopoietic stem cells in vivo using high-throughput sequencing in conjunction with viral genetic barcoding. Nat. Biotechnol. 29, 928–933. 10.1038/nbt.1977.

8. Naik, S.H., Perié, L., Swart, E., Gerlach, C., van Rooij, N., de Boer, R.J., and Schumacher, T.N. (2013). Diverse and heritable lineage imprinting of early haematopoietic progenitors. Nature 496, 229–232. 10.1038/nature12013.

9. Rehman, S.K., Haynes, J., Collignon, E., Brown, K.R., Wang, Y., Nixon, A.M.L., Bruce, J.P., Wintersinger, J.A., Singh Mer, A., Lo, E.B.L., et al. (2021). Colorectal Cancer Cells Enter a Diapause-like DTP State to Survive Chemotherapy. Cell 184, 226–242.e21. 10.1016/j.cell.2020.11.018.

10. Gutierrez, C., Al’Khafaji, A.M., Brenner, E., Johnson, K.E., Gohil, S.H., Lin, Z., Knisbacher, B.A., Durrett, R.E., Li, S., Parvin, S., et al. (2021). Multifunctional barcoding with ClonMapper enables high-resolution study of clonal dynamics during tumor evolution and treatment. Nat. Cancer 2, 758–772. 10.1038/s43018-021-00222-8.

11. Oren, Y., Tsabar, M., Cuoco, M.S., Amir-Zilberstein, L., Cabanos, H.F., Hütter, J.-C., Hu, B., Thakore, P.I., Tabaka, M., Fulco, C.P., et al. (2021). Cycling cancer persister cells arise from lineages with distinct programs. Nature 596, 576–582. 10.1038/s41586-021-03796-6.

12. Scherer, M., Singh, I., Braun, M.M., Szu-Tu, C., Sanchez Sanchez, P., Lindenhofer, D., Jakobsen, N.A., Körber, V., Kardorff, M., Nitsch, L., et al. (2025). Clonal tracing with somatic epimutations reveals dynamics of blood ageing. Nature 643, 478–487. 10.1038/s41586-025-09041-8.

13. Rodriguez-Fraticelli, A.E., Weinreb, C., Wang, S.-W., Migueles, R.P., Jankovic, M., Usart, M., Klein, A.M., Lowell, S., and Camargo, F.D. (2020). Single-cell lineage tracing unveils a role for TCF15 in haematopoiesis. Nature 583, 585–589. 10.1038/s41586-020-2503-6.

14. He, J., Qiu, Z., Fan, J., Xie, X., Sheng, Q., and Sui, X. (2024). Drug tolerant persister cell plasticity in cancer: A revolutionary strategy for more effective anticancer therapies. Signal Transduct. Target. Ther. 9, 209. 10.1038/s41392-024-01891-4.

15. Sharma, S.V., Lee, D.Y., Li, B., Quinlan, M.P., Takahashi, F., Maheswaran, S., McDermott, U., Azizian, N., Zou, L., Fischbach, M.A., et al. (2010). A chromatin-mediated reversible drug-tolerant state in cancer cell subpopulations. Cell 141, 69–80. 10.1016/j.cell.2010.02.027.

16. Jones, T.P., and McGranahan, N. (2023). Deciphering the landscape of transcriptional heterogeneity across cancer. Cancer Cell 41, 1548–1550. 10.1016/j.ccell.2023.07.008.

17. Feinberg, A.P., and Levchenko, A. (2023). Epigenetics as a mediator of plasticity in cancer. Science 379, eaaw3835. 10.1126/science.aaw3835.

18. Gupta, P.B., Fillmore, C.M., Jiang, G., Shapira, S.D., Tao, K., Kuperwasser, C., and Lander, E.S. (2011). Stochastic state transitions give rise to phenotypic equilibrium in populations of cancer cells. Cell 146, 633–644. 10.1016/j.cell.2011.07.026.

19. Beerling, E., Seinstra, D., de Wit, E., Kester, L., van der Velden, D., Maynard, C., Schäfer, R., van Diest, P., Voest, E., van Oudenaarden, A., et al. (2016). Plasticity between Epithelial and Mesenchymal States Unlinks EMT from Metastasis-Enhancing Stem Cell Capacity. Cell Rep. 14, 2281–2288. 10.1016/j.celrep.2016.02.034.

20. Pastushenko, I., Brisebarre, A., Sifrim, A., Fioramonti, M., Revenco, T., Boumahdi, S., Van Keymeulen, A., Brown, D., Moers, V., Lemaire, S., et al. (2018). Identification of the tumour transition states occurring during EMT. Nature 556, 463–468. 10.1038/s41586-018-0040-3.

21. Nguyen, L.V., Eyal-Lubling, Y., Guerrero-Romero, D., Kronheim, S., Chin, S.-F., Manzano Garcia, R., Sammut, S.-J., Lerda, G., Lui, A.J.W., Bardwell, H.A., et al. (2025). Fitness and transcriptional plasticity of human breast cancer single-cell-derived clones. Cell Rep. 44, 115699. 10.1016/j.celrep.2025.115699.

22. Alonso, S., Raghav, K., Morris, V.K., Alfaro-Munoz, K., Bekaii-Saab, T., Cannon, T.L., Corcoran, R.B., Duesbery, N., George, M., Hsu, D., et al. (2026). Framework for cancer evolution profiling and interception in colorectal cancer: ASCEND-CRC program. Cancer Cell 44, 455–459. 10.1016/j.ccell.2025.12.016.

23. Niture, S., Mooers, B.H.M., Wu, D.H., Hart, M., Jaboin, J., and Seneviratne, D. (2025). Dual-specificity protein phosphatase 1: A potential therapeutic target in cancer. iScience 28, 113706. 10.1016/j.isci.2025.113706.

24. Shen, J., Zhang, Y., Yu, H., Shen, B., Liang, Y., Jin, R., Liu, X., Shi, L., and Cai, X. (2016). Role of DUSP1/MKP1 in tumorigenesis, tumor progression and therapy. Cancer Med. 5, 2061–2068. 10.1002/cam4.772.

25. Wang, J., Kho, D.H., Zhou, J.-Y., Davis, R.J., and Wu, G.S. (2017). MKP-1 suppresses PARP-1 degradation to mediate cisplatin resistance. Oncogene 36, 5939–5947. 10.1038/onc.2017.197.

26. Sanders, B.E., Yamamoto, T.M., McMellen, A., Woodruff, E.R., Berning, A., Post, M.D., and Bitler, B.G. (2022). Targeting DUSP Activity as a Treatment for High-Grade Serous Ovarian Carcinoma. Mol. Cancer Ther. 21, 1285–1295. 10.1158/1535-7163.MCT-21-0682.

27. Yadav, S.S., Kalia, P., Kaur, N., Sharma, S., Thakur, S., Kumari, S., and Nair, R.R. (2025). KLF4 in cancer chemoresistance: molecular mechanisms and therapeutic implications. Discov. Oncol. 16, 1690. 10.1007/s12672-025-02261-4.

28. He, Z., He, J., and Xie, K. (2023). KLF4 transcription factor in tumorigenesis. Cell Death Discov. 9, 118. 10.1038/s41420-023-01416-y.

29. Magbanua, M.J.M., Wolf, D.M., Yau, C., Davis, S.E., Crothers, J., Au, A., Haqq, C.M., Livasy, C., Rugo, H.S., I-SPY 1 TRIAL Investigators, et al. (2015). Serial expression analysis of breast tumors during neoadjuvant chemotherapy reveals changes in cell cycle and immune pathways associated with recurrence and response. Breast Cancer Res. BCR 17, 73. 10.1186/s13058-015-0582-3.

30. Bhang, H.C., Ruddy, D.A., Krishnamurthy Radhakrishna, V., Caushi, J.X., Zhao, R., Hims, M.M., Singh, A.P., Kao, I., Rakiec, D., Shaw, P., et al. (2015). Studying clonal dynamics in response to cancer therapy using high-complexity barcoding. Nat. Med. 21, 440–448. 10.1038/nm.3841.

31. Emert, B.L., Cote, C.J., Torre, E.A., Dardani, I.P., Jiang, C.L., Jain, N., Shaffer, S.M., and Raj, A. (2021). Variability within rare cell states enables multiple paths toward drug resistance. Nat. Biotechnol. 39, 865–876. 10.1038/s41587-021-00837-3.

32. Shea, A., Eyal-Lubling, Y., Guerrero-Romero, D., Manzano Garcia, R., Greenwood, W., O’Reilly, M., Georgopoulou, D., Callari, M., Lerda, G., Wix, S., et al. (2025). Modeling Drug Responses and Evolutionary Dynamics Using Patient-Derived Xenografts Reveals Precision Medicine Strategies for Triple-Negative Breast Cancer. Cancer Res. 85, 567–584. 10.1158/0008-5472.CAN-24-1703.

33. Rodriguez-Fraticelli, A.E., Weinreb, C., Wang, S.-W., Migueles, R.P., Jankovic, M., Usart, M., Klein, A.M., Lowell, S., and Camargo, F.D. (2020). Single-cell lineage tracing unveils a role for TCF15 in haematopoiesis. Nature 583, 585–589. 10.1038/s41586-020-2503-6.

34. Bruna, A., Rueda, O.M., Greenwood, W., Batra, A.S., Callari, M., Batra, R.N., Pogrebniak, K., Sandoval, J., Cassidy, J.W., Tufegdzic-Vidakovic, A., et al. (2016). A Biobank of Breast Cancer Explants with Preserved Intra-tumor Heterogeneity to Screen Anticancer Compounds. Cell 167, 260–274.e22. 10.1016/j.cell.2016.08.041.

35. Shin, H.J., Sharma, S., Kronheim, S., Cheung, M., and Nguyen, L.V. (2025). Protocol to track single-cell-derived clones using DNA barcoding combined with single-cell RNA sequencing. STAR Protoc. 6, 104229. 10.1016/j.xpro.2025.104229.

36. Zheng, G.X.Y., Terry, J.M., Belgrader, P., Ryvkin, P., Bent, Z.W., Wilson, R., Ziraldo, S.B., Wheeler, T.D., McDermott, G.P., Zhu, J., et al. (2017). Massively parallel digital transcriptional profiling of single cells. Nat. Commun. 8, 14049. 10.1038/ncomms14049.

37. Hao, Y., Stuart, T., Kowalski, M.H., Choudhary, S., Hoffman, P., Hartman, A., Srivastava, A., Molla, G., Madad, S., Fernandez-Granda, C., et al. (2024). Dictionary learning for integrative, multimodal and scalable single-cell analysis. Nat. Biotechnol. 42, 293–304. 10.1038/s41587-023-01767-y.

38. Germain, P.-L., Lun, A., Garcia Meixide, C., Macnair, W., and Robinson, M.D. (2022). Doublet identification in single-cell sequencing data using scDblFinder. F1000Research 10, 979. 10.12688/f1000research.73600.2.

39. Zappia, L., and Oshlack, A. (2018). Clustering trees: a visualization for evaluating clusterings at multiple resolutions. GigaScience 7, giy083. 10.1093/gigascience/giy083.

40. Wickham, H. (2016). ggplot2: elegant graphics for data analysis Second edition. (Springer international publishing).

41. Blighe, K. (2018). EnhancedVolcano. (Bioconductor). 10.18129/B9.BIOC.ENHANCEDVOLCANO https://doi.org/10.18129/B9.BIOC.ENHANCEDVOLCANO.

42. The Gene Ontology Consortium, Aleksander, S.A., Balhoff, J., Carbon, S., Cherry, J.M., Drabkin, H.J., Ebert, D., Feuermann, M., Gaudet, P., Harris, N.L., et al. (2023). The Gene Ontology knowledgebase in 2023. GENETICS 224, iyad031. 10.1093/genetics/iyad031.

43. Ashburner, M., Ball, C.A., Blake, J.A., Botstein, D., Butler, H., Cherry, J.M., Davis, A.P., Dolinski, K., Dwight, S.S., Eppig, J.T., et al. (2000). Gene Ontology: tool for the unification of biology. Nat. Genet. 25, 25–29. 10.1038/75556.

44. Yu, G., Wang, L.-G., Han, Y., and He, Q.-Y. (2012). clusterProfiler: an R Package for Comparing Biological Themes Among Gene Clusters. OMICS J. Integr. Biol. 16, 284–287. 10.1089/omi.2011.0118.

45. Yu, G., Li, F., Qin, Y., Bo, X., Wu, Y., and Wang, S. (2010). GOSemSim: an R package for measuring semantic similarity among GO terms and gene products. Bioinformatics 26, 976–978. 10.1093/bioinformatics/btq064.

46. Guangchuang Yu (2018). enrichplot. (Bioconductor). 10.18129/B9.BIOC.ENRICHPLOT https://doi.org/10.18129/B9.BIOC.ENRICHPLOT.

47. Gu, Z., and Hübschmann, D. (2023). *SimplifyEnrichment*: A Bioconductor Package for Clustering and Visualizing Functional Enrichment Results. Genomics Proteomics Bioinformatics 21, 190–202. 10.1016/j.gpb.2022.04.008.

48. Andreatta, M., and Carmona, S.J. (2025). GeneNMF: Non-Negative Matrix Factorization for Single-Cell Omics. Version 0.9.2.

49. Andreatta, M., and Carmona, S.J. (2021). UCell: Robust and scalable single-cell gene signature scoring. Comput. Struct. Biotechnol. J. 19, 3796–3798. 10.1016/j.csbj.2021.06.043.

